# Colour and shape evolution reflect ecological specialisation in Pomacentridae

**DOI:** 10.1101/2025.08.05.668455

**Authors:** Alberto García Jiménez, Nicolas Salamin, Théo Gaboriau

## Abstract

Damselfishes (Pomacentridae) display remarkable diversity in colouration and body form, yet the processes shaping this phenotypic variation remain poorly resolved. Our study aimed to characterise the evolution of these traits, evaluate their associations with ecological factors, and identify convergent patterns linked to ecological specialisation. Using image-based quantification of colour patterns, geometric morphometrics, and phylogenetic comparative methods across 343 species, we show that pomacentrid phenotypes are organised around a small number of dominant axes describing brightness, hue, contrast, body elongation, and cranial morphology. Both colour and morphology exhibit early bursts of evolutionary disparity, followed by recurrent lineage-specific radiations and widespread convergence toward similar adaptive optima across the phylogeny. Dietary ecotypes emerged as the strongest predictor of morphological diversification, whereas symbiotic and social regimes exerted the strongest effects on colour evolution. Despite these distinct ecological correlates, several colour and shape axes form partially integrated trait syndromes that evolve in concert. The pervasive convergence of colour–shape syndromes underscores deterministic components of reef-fish evolution and positions Pomacentridae as a model for understanding integrated phenotypic evolution.

## Introduction

Understanding how ecological pressures shape phenotypic evolution is a central goal of evolutionary biology. In reef fishes, two major axes of phenotypic variation —body morphology and colouration— play key roles in mediating ecological performance, including locomotion, feeding, camouflage, and social signalling [1–3]. Morphological traits are tightly linked to habitat use and trophic strategies, and their macroevolutionary dynamics are increasingly well understood, often revealing strong convergence driven by repeated ecological transitions across reef environments [4–6]. By contrast, the evolutionary dynamics of colouration, and its relationship to morphology, remain comparatively poorly resolved at macroevolutionary scales.

Colour patterns in reef fishes mediate diverse and context-dependent functions, including crypsis, communication, mate choice, and species recognition, shaped by visual environments and behavioural interactions [3, 7]. Despite this functional diversity, comparative studies of colour evolution have frequently relied on simplified or discrete categorizations of patterns, such as stripes, bars, or eyespots [8, 9]. While informative, these approaches may obscure continuous and multivariate variation in pigmentation that can encode ecologically relevant information [10, 11]. Recent methodological advances now enable quantitative, image-based analyses of colour patterns as continuous traits, opening new opportunities to examine colour evolution alongside morphology within a unified comparative framework [12, 13].

Integrating colour and morphology is particularly important because these traits may not evolve independently. Colouration can enhance or modulate the ecological functions of body form, for example by reinforcing camouflage in specific habitats or amplifying visual signals associated with social or symbiotic behaviours [14, 15]. However, whether colour and morphology evolve largely as independent axes responding to distinct ecological drivers, or as partially integrated trait suites shaped by shared selective pressures, remains largely unexplored across species-rich reef-fish clades.

The damselfishes and clownfishes (Pomacentridae) provide an exceptional system to address these questions. With over 400 species distributed across a wide range of habitats, depths, and trophic strategies, pomacentrids rank among the most diverse and ecologically important reef-fish families [16, 17]. They exhibit striking diversity in colouration, ranging from uniform or gradient-based pigmentation to highly contrasting patterns involving bars, bands, and localized markings [18], alongside substantial variation in body form, fin proportions, and cranial morphology [2, 5]. Pomacentrids span a continuum from pelagic planktivores to benthic algal farmers and include both free-living species and lineages engaged in commensal or mutualistic interactions [19, 20]. These ecological dimensions tend to covary, forming recurrent ecological syndromes or “ecotypes” that integrate feeding mode, microhabitat specialization, and behaviour, and provide a framework for linking joint ecological constraints to phenotypic evolution across the family [2, 6].

Previous comparative work has shown that pomacentrid morphology exhibits strong convergence across these ecotypes, with repeated transitions driving iterative evolution of jaw structure, body depth, and locomotor form [4, 21]. In contrast, evidence for predictable or convergent colour evolution remains limited, largely restricted to clownfishes and their anemone-associated niche [22]. This raises a broader question: is colour convergence a unique feature of clownfishes, or, like morphological traits, does it reflect repeated evolutionary responses to shared ecological pressures across pomacentrids? Importantly, considering morphology and colour jointly, rather than separately, may be essential to understand whether ecological diversification within pomacentrids is driven by coordinated or independent evolutionary responses.

We test these questions using a comparative framework integrating quantitative colour-pattern analyses, geometric morphometrics, ecological information and phylogenetic comparative methods across 343 species of Pomacentrids (315 damselfishes and 28 clownfishes). Using multivariate trait spaces and explicit ecological predictors, we test how diet, habitat, and symbiotic behaviour shape the evolution of colour and morphology, assess the prevalence of convergence across both trait domains, and quantify the extent of evolutionary integration between them. By treating colour and body shape as coupled components of a broader phenotypic system, our study provides a unified macroevolutionary perspective on how complex phenotypes diversify in structured reef environments.

## Results

### Patterns of colour and morphological diversity

Principal component analysis of 1,183 Procrustes-aligned colour-pattern images from 343 pomacentrid species revealed a low-dimensional structure, with the first eight PCs explaining 48.8% of total colour variation. PC1 (17.8%) captured a gradient from light to dark pigmentation, PC2 (11.5%) separated reddish from bluish hues and rostral white bar presence, and PC3–PC4 described body– edge and dorsoventral patterning (Figure 2a,b). Higher PCs (PC5–PC8) captured more localized pattern elements, including segmentation and contrast variation (Supplementary Figure S2a,b).

**Fig. 1:**
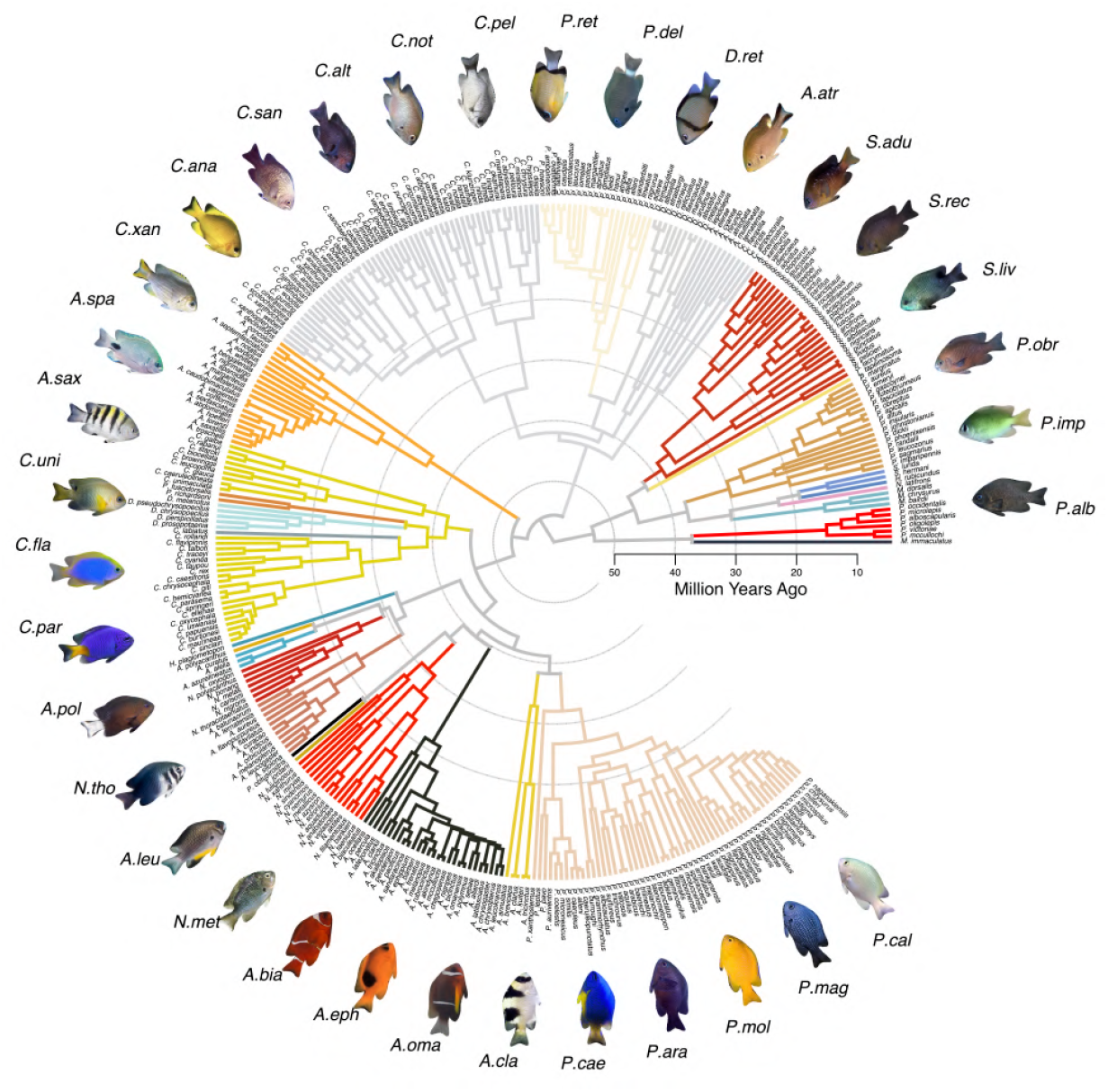
Circular phylogenetic tree sourced from McCord *et al*. (2021), with branches colour-coded to distinguish species by genus. Procrustes-transformed images of 34 out of the 342 species utilized in this study are displayed in the outer margin encircling the tree. Species names are abbreviated using the initial of the genus followed by the first three characters of the species name. Horizontal scale depicts time in million years before the present (Mya).

**Fig. 2:**
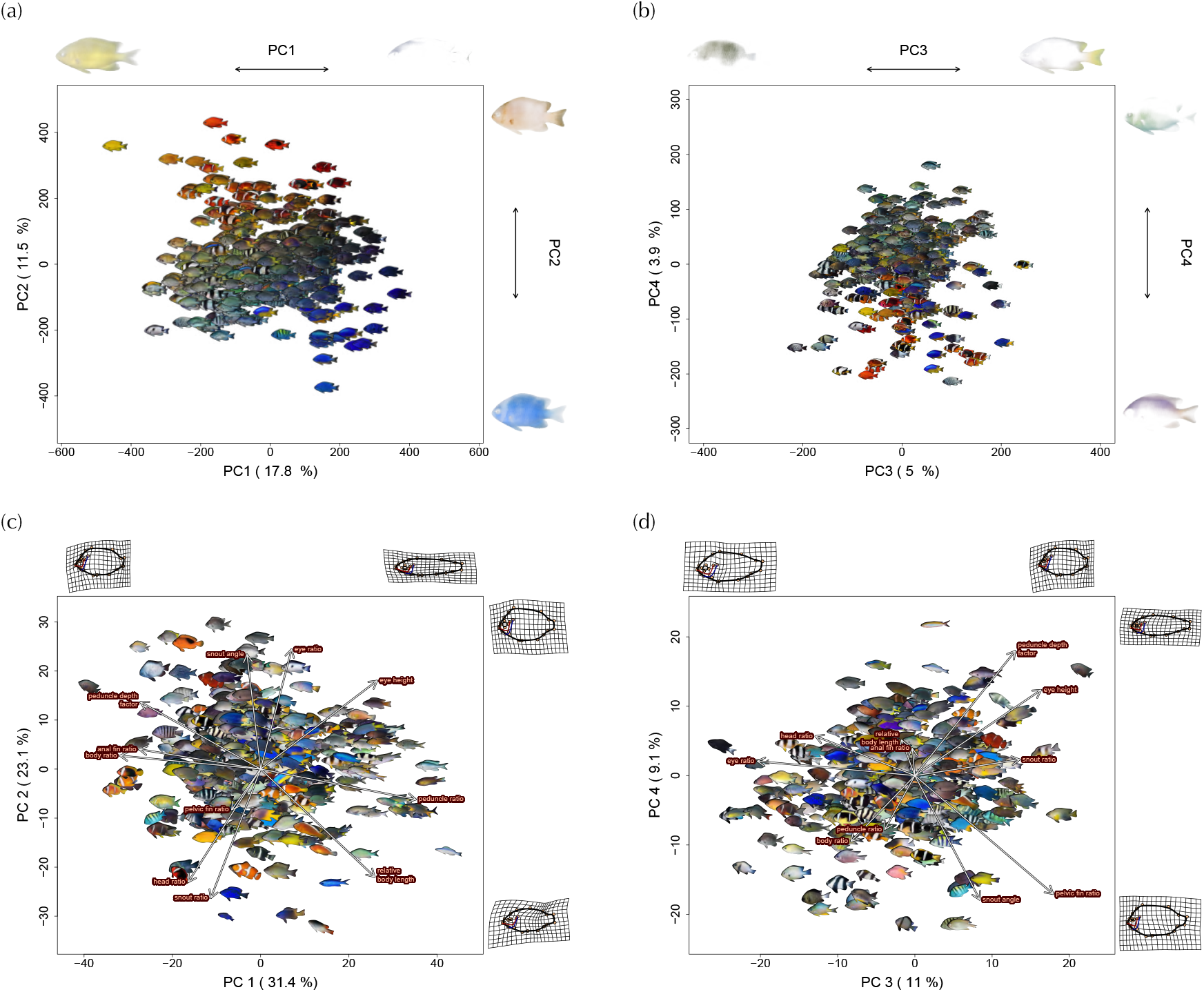
Principal component analyses of colour patterns and body shape across 343 pomacentrid species. (a,b) Colourpattern PCA based on 1,183 Procrustes-aligned images. (c,d) Morphological PCA based on 11 morphometric traits. Axis labels indicate principal component and variance explained. In the colour PCA, points represent species-averaged pixel coordinates, with reconstructed mean images projected onto the morphospace; marginal images illustrate extreme colour-pattern values, with pixel opacity scaled by axis contribution. In the morphology PCA, background-removed specimen images are shown, with eigenvector loadings indicated by white arrows and marginal deformation grids illustrating shape extremes.

Morphological PCA of 11 standardized traits similarly revealed a compact phenotypic space, with the first eight PCs accounting for 96.1% of total variation (Figure 2c,d). PC1 (31.4%) described a depth–elongation gradient, PC2 (23.1%) contrasted head and snout proportions, and subsequent PCs captured variation in eye position, peduncle depth, and fin proportions relevant to locomotion and feeding (Supplementary Figure S2c,d).

### Phenotypic variance and diversification dynamics

PERMANOVA revealed strong species-level structuring of phenotypic variation, with species identity explaining nearly 70% of total individual-level colour and morphological variance (*p* = 0.001). In contrast, broad ecological regimes explained modest proportions of species-averaged variation: diet (*R*^2^ = 0.0286, *F* = 5.38, *p* = 0.001), habitat (*R*^2^ = 0.0215, *F* = 2.69, *p* = 0.001), and symbiosis (*R*^2^ = 0.0033, *F* = 1.242, *p* = 0.236), with residual variation exceeding 87%.

Disparity-through-time (DTT) analyses revealed three recurring modes of phenotypic diversification across colour and morphological axes (Figure 3).

**Fig. 3:**
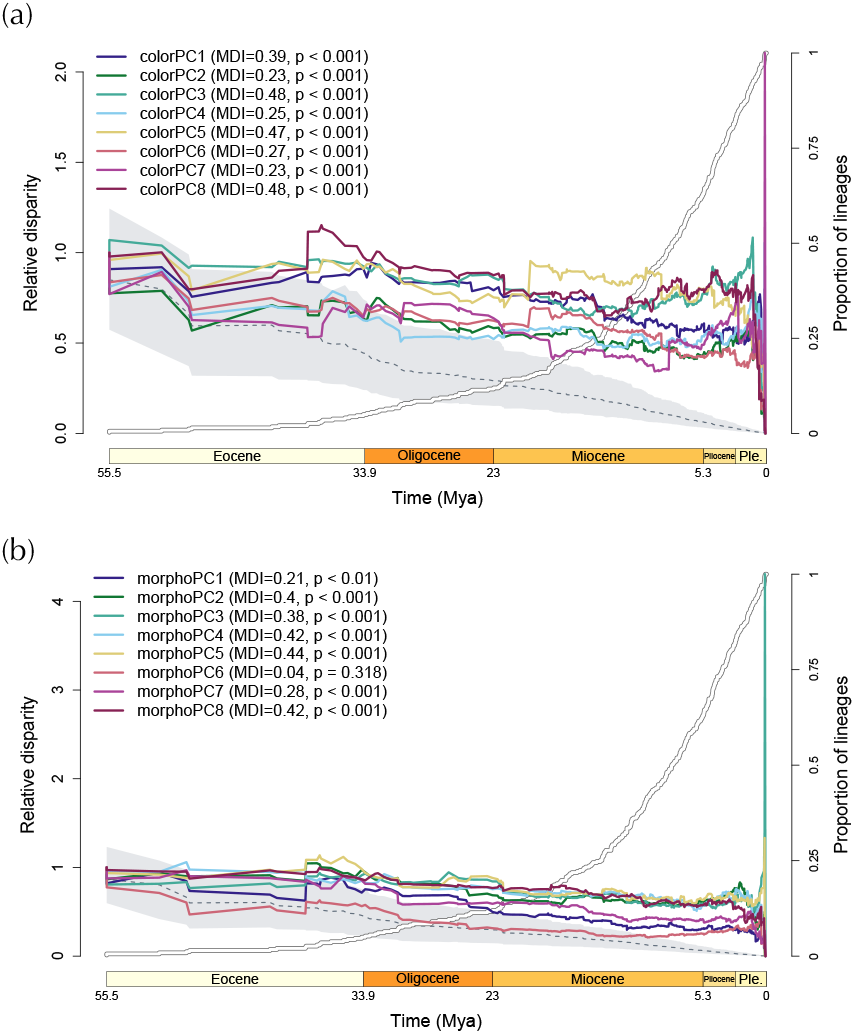
DTT plots for colour (a) and morphology (b) PCs 1–8. Coloured lines show observed disparity per PC, thick white lines the LTT curve, and dashed grey with shading the BM expectation (95% CI). MDI and *p*-values at top quantify deviation from BM.

#### (1) Early-burst trajectories

Colour PC1, PC3, PC5, and PC8, together with morphology PC2–PC5 and PC8, exhibited strong early bursts of disparity, with pronounced peaks around ∼ 40 Mya (relative time ∼ 0.3). These axes showed high positive MDI values (0.30–0.48) and significant deviation from Brownian motion (*p* < 0.01), indicating rapid early partitioning of phenotypic space among major clades. They capture major gradients in colour contrast, body–edge pigmentation, body depth, fin proportions, and peduncle morphology. Following this early diversification, disparity trajectories converged toward Brownian expectations, consistent with stabilisation of among-clade differences and subsequent within-clade diversification.

#### (2) Gradual-accumulation trajectories

Colour PC2, PC4, PC6, and PC7, together with morphology PC1 and PC7, showed progressive increases in disparity through time (MDI = 0.20–0.30), reflecting steady within-clade diversification rather than early divergence. These axes describe variation in hue, dorsoventral or lateral segmentation, overall body elongation, and head–snout morphology. Several PCs also exhibited secondary late increases in disparity near the present (relative time > 0.8), indicating recent diversification within younger subclades.

#### (3) Brownian trajectories

Morphology PC6 closely followed Brownian expectations (MDI ≈ 0, *p* = 0.273), consistent with neutral or weakly constrained evolution across most of the tree. Nonetheless, a modest increase in disparity among recent lineages suggests limited late-stage expansion of phenotypic space.

### Regime shifts and convergence in colour and morphology

Across colour and morphological trait space, *ℓ*1ou detected widespread shifts in adaptive optima, with repeated transitions occurring across multiple lineages and trait axes (Figure 4 and Supplementary Tables S1–S2). Well-supported shifts (bootstrap support > 0.7) were especially frequent on colour components associated with contrast and patterning (PC2, PC4, PC6, PC7), which capture major elements such as white bars, dorsoventral contrast, and lateral pigmentation.

**Fig. 4:**
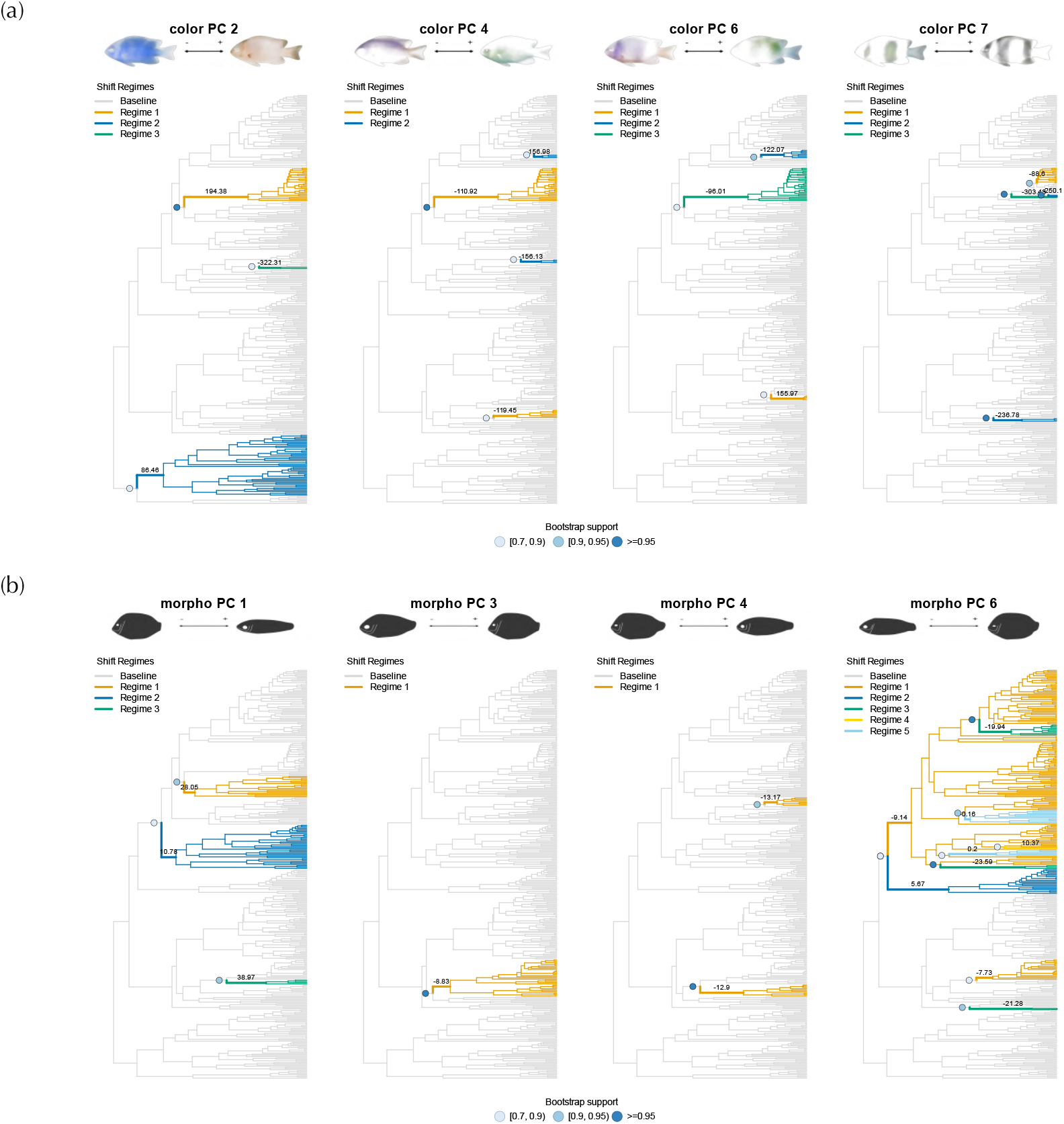
*ℓ*1ou-inferred regime shifts across the damselfish phylogeny for biologically relevant colour (a) and morphology (b) PCs (bs > 0.7). Coloured branches show convergent shifts to alternative optima (grey: ancestral); filled circles indicate bootstrap support. Reconstructed patterns/deformations illustrate PC extremes. Full results in Supplementary Figs. S3–S4.

Clownfishes (*Amphiprion*) showed multiple transitions toward extreme colour optima, including high-contrast patterns on PC2 and darker optima on PC4 and PC6, with additional lineage-specific shifts in the skunk complex. Comparable contrast-reducing or darkening shifts occurred independently in *Chrysiptera, Dascyllus*, and *Pomacentrus*, indicating recurrent evolution of similar pigmentation syndromes across reef-associated lineages. Lineage-specific shifts with moderate support included strong darkening in *Hypsypops* (PC4, PC8) and brightening in *Stegastes partitus* and *Chromis vanbebberae* (PC5).

Morphological evolution displayed similarly pervasive convergence. Axes describing body elongation and depth (PC1–PC5) showed repeated transitions across the phylogeny, with multiple lineages independently evolving elongated, mid-water morphologies (e.g. *Chromis, Azurina, Neopomacentrus, Pristotis*, and *Teixeirichthys*) or deeper-bodied forms associated with benthic or symbiotic lifestyles (e.g. *Chrysiptera, Cheiloprion, Amphiprion*, and *Pomacentrus*). Additional convergent shifts were detected along axes describing head shape and compression axes (PC6–PC8), including repeated transitions toward fusiform profiles in pelagic taxa and more compact cranial morphologies in reef-associated clades.

Across the phylogeny, shifts in colour and morphology frequently co-occurred. Lineages such as *Amphiprion, Chrysiptera*, and *Dascyllus* combined transitions toward high-contrast or darker colour patterns with changes in body depth and cranial morphology, whereas pelagic-oriented taxa such as *Chromis* and *Azurina* consistently coupled elongated body forms with low-contrast or gradient-based colour axes.

### Comparative analyses of colour and morphology under ecological regimes

Across colour and morphological datasets, Pagel’s *λ* provided the best-fitting phylogenetic correlation structure for most axes, indicating moderate phylogenetic dependence (Supplementary Tables S3–S6). Ecological predictors consistently outperformed null models in multivariate analyses, whereas univariate models more often favoured intercept-only fits, with significant ecological effects restricted to a subset of PC axes (Supplementary Tables S7, S8). Ancestral trait reconstructions used to inform these analyses are reported in Supplementary Results.

Diet was the strongest predictor of joint colour– morphology variation in multivariate PGLS (EIC = 48,701.525; *EIC*_*w*_ = 0.86), supported by PERMANOVA (Pillai = 0.225, *p* = 0.001) and moderate phylogenetic signal (*λ* = 0.50; Supplementary Table S9). Diet also best explained morphological variation alone (EIC = 22,159.287; *EIC*_*w*_ = 1.00; *λ* = 0.58; Pillai = 0.177, *p* = 0.001; Supplementary Table S10). Relative to benthic feeders, intermediate-diet species showed reduced brightness and bluish hues (colPC1, colPC2), rostral markings (colPC6), and increased ventral–caudal brightness (colPC4–5, colPC8), alongside shifts toward more elongated bodies with taller peduncles and larger eyes (morphoPC1–2). Pelagic feeders exhibited stronger colour shifts toward light-blue and silvery phenotypes and pronounced body elongation (morphoPC1) compared to benthic taxa.

Symbiosis received secondary support in the combined analysis (*EIC*_*w*_ = 0.14) but was the best predictor of colour variation alone (EIC = 31,043.720; *EIC*_*w*_ = 1.00; *λ* = 0.48; *Pillai* = 0.112, *p* = 0.005; Supplementary Table S11). Effect sizes mirrored those from colour-only models: commensalistic species showed positive shifts on colPC1 and colPC2, while mutualistic taxa exhibited strong positive effects on colPC2 and negative shifts on axes describing body-edge contrast, dorsoventral gradients, and lateral segmentation (colPC3, colPC4, colPC6).

Multivariate trait covariances revealed consistent axes of integration between colour and morphology (Supplementary Figure S5). Light–dark variation (colPC1) covaried negatively with body shape, head–eye geometry, and ventral/anal fin configurations (morphoPC1–2, morphoPC5– 6), whereas ventral–caudal brightness (colPC4–5) covaried positively with elongated, high-peduncle body forms (morphoPC1). Bluish–reddish hue and rostral markings (colPC2) were negatively associated with body elongation, and pattern segmentation axes (colPC3, colPC7) showed strong covariation with head and eye morphology (morphoPC2–3), indicating coordinated evolution of pigmentation and cranial morphology.

### Model comparison of colour and morphological trait evolution

Comparative analyses of trait evolutionary models consistently rejected simple models like BM or WN in favour of more complex processes (Supplementary Figure S6a). For both colour and morphology, multi-optimum OU models (OUM) were most frequently preferred, often with strong support (high Akaike weights and strong ERs), while diversity-dependent (DD) models were occasionally selected (Supplementary Figure S7a–b). Four morphological axes (PC1, PC3, PC6 and PC7) were best explained by an exponential density-dependent model (DDexp), whereas the remaining morphological and all colour axes were consistently supported by OUM models structured by ecological regimes (diet, habitat or symbiosis). DDexp-supported axes showed uniformly positive diversity-dependence parameters (PC1: 0.012; PC3: 0.137; PC6: 0.012; PC7: 0.015), indicating accelerating evolutionary rates with increasing lineage diversity. PC3 exhibited the strongest effect, while the rest showed weaker signals with limited stochastic diffusion (*σ*^2^ ≈ 0–0.004).

### Integration between colour and morphology

Univariate PGLS analyses revealed six significant correlations between morphological and colour axes after FDR correction (Figure 5 and Supplementary Table S12). MorphoPC1, reflecting body elongation, positively covaried with colorPC5 (*β* = 0.167, *p*_*FDR*_ = 0.035; Pagel’s *λ*), which captures antero-posterior pigmentation gradients. MorphoPC2, primarily describing head shape, was positively associated with colorPC3 (*β* = 0.158, *p*_*FDR*_ = 0.040; Pagel’s *λ*), contrasting body and edge pigmentation. MorphoPC3 (eye–snout covariation) positively correlated with colorPC7 (*β* = 0.177, *p*_*FDR*_ = 0.013; Pagel’s *λ*), representing vertical barring patterns. MorphoPC4 (snout and peduncle depth) was negatively linked to colorPC6 (*β* = − 0.161, *p*_*FDR*_ = 0.027; Pagel’s *λ*), distinguishing anteroventral from posterodorsal colour contrasts. Finally, morphoPC6 (body and anal fin proportions) showed inverse associations with both colorPC5 (*β* = − 0.143, *p*_*FDR*_ = 0.035; OU) and colorPC7 (*β* = − 0.185, *p*_*FDR*_ = 0.046; Pagel’s *λ*).

**Fig. 5:**
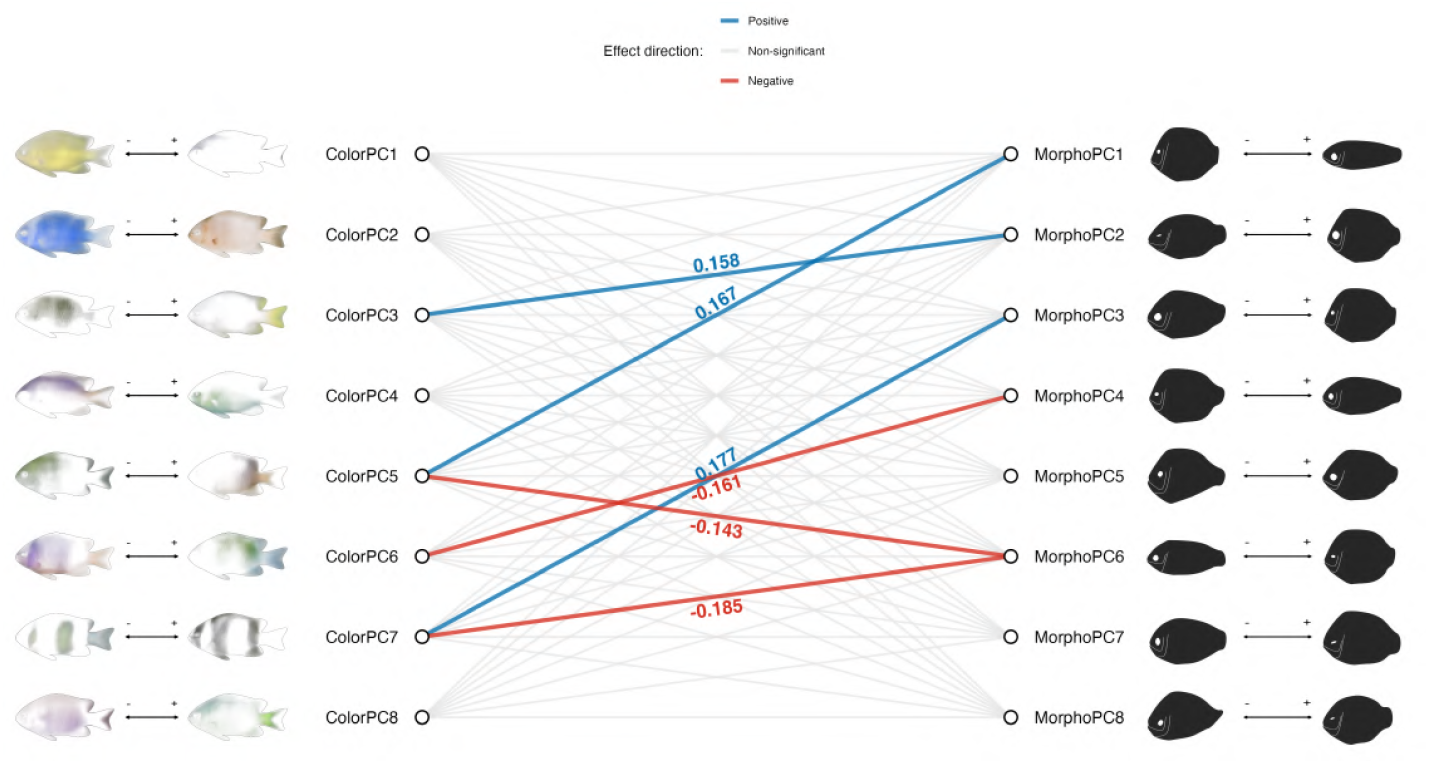
Colour–morphology PC integration. Left/right panels show reconstructed phenotypic extremes and pixel contributions for colour/morphology PCs. Central network displays significant PGLS associations (FDR *p* < 0.05; red/blue edges: positive/negative *β*).

Multivariate PGLS identified the trait model with morphology under a Pagel’s *λ* correlation structure (EIC = 31,051.47; weight = 0.995) as a better fit than the null model (ΔEIC = 10.769; weight = 0.456; Supplementary Table S13). PERMANOVA confirmed a significant association (Pillai’s trace = 0.356, *p* = 0.001), indicating that colour and morphological traits are evolutionarily correlated and exhibit partial integration across the Pomacentridae phylogeny.

## Discussion

Pomacentrid diversification has produced exceptional diversity in colouration, body form, and ecology, yet the processes structuring this phenotypic radiation have remained unclear. By integrating high-resolution colour-pattern data with morphometrics and phylogenetic comparative analyses across 343 species, we show that pomacentrid phenotypes evolve along a small number of dominant axes, experience rapid early diversification followed by recurrent lineage-specific radiations, and repeatedly converge on similar adaptive optima. Importantly, colour and body shape are not independent dimensions, but form partially integrated phenotypic syndromes shaped by shared ecological pressures. Our results provide a unified view of how ecological regimes, diversification dynamics, and adaptive landscapes interact to generate the characteristic visual and morphological diversity of Pomacentridae.

### Mapping the phenotypic space of pomacentrids

Colour variation in pomacentrids is structured primarily along axes describing brightness, red–blue chromatic balance, and contrast or segmentation. The light–dark gradient represents a fundamental dimension of reef-fish pigmentation, mediating both camouflage and visual signalling across heterogeneous light environments, while yellow–blue balance reflects perceptual and behavioural processes linked to habitat filtering, mate choice, and social recognition [3]. These continuous dimensions would be poorly captured by discrete scoring schemes, which cannot resolve gradual shifts, combined effects, or integration among colour traits. Consistent with multivariate analyses in other reef fishes [11, 12], the colour axes recovered here structure major ecological associations and recurrent evolutionary trajectories, demonstrating that continuous colour spaces are essential for capturing functionally and evolutionarily meaningful pigmentation variation across reef-fish clades [13].

Morphological variation resolves into similarly interpretable ecological gradients. The primary morphological axis captures an elongation–depth continuum separating streamlined pelagic planktivores from deeper-bodied benthic, territorial, or symbiotic taxa, mirroring a dominant body–shape gradient observed across Indo-Pacific reef fishes [5]. A second axis reflects variation in head and snout proportions associated with feeding mode, while higher-order axes describe modular fin and cranial adjustments that fine-tune locomotion and feeding performance, mirroring recurrent functional trade-offs underlying ecological specialization in reef fishes [4, 21].

### Adaptive diversification and ecological convergence

Disparity–through–time analyses reveal rapid early diversification of both colour and morphology, as early pomacentrid lineages partitioned ecological and phenotypic space and established the main axes of variation. This pattern mirrors other reef-fish radiations associated with ecological opportunity during Miocene reef expansion [23, 24]. However, disparity did not remain canalised, revealing multiple late bursts occurred within derived clades, consistent with repeated transitions among ecological strategies such as pelagic versus benthic feeding or renewed reef colonisation [25, 26]. These pulses indicate that evolutionary innovation persisted long after initial niche filling, highlighting the iterative nature of pomacentrid diversification.

Consistent with this dynamic, *ℓ*1ou analyses show that many late increases in disparity reflect convergence toward shared adaptive optima. Phylogenetically distant lineages repeatedly evolved similar pigmentation gradients and body forms, occupying comparable regions of phenotypic space despite independent origins. This pervasive convergence reinforces that pomacentrids diversify within a constrained set of viable ecomorphological solutions imposed by reef environments [4]. Recurrent colour patterns in symbiotic and territorial taxa, together with repeated evolution of deep-bodied herbivores and elongate planktivores, reflect parallel adaptive responses to similar ecological contexts, reinforcing Pomacentridae as an archetypal example of iterative ecological radiation.

### Ecology-specific drivers of colour and morphology

Although phylogeny explained much of the variance in colour and shape, ecological effects were detectable and trait-specific. Diet emerged as the strongest predictor of morphological evolution, supporting extensive evidence that trophic diversification underpins reef-fish functional evolution [6, 27]. The repeated emergence of deep-bodied algal farmers and streamlined planktivores parallels patterns observed in other herbivorous and planktivorous reef-fish clades, where feeding mode imposes consistent biomechanical and cranial constraints [21, 28].

In contrast, colour evolution was most strongly associated with symbiotic and social regimes. Pronounced colour divergence in anemone-associated lineages supports the role of visually mediated interactions with hosts and conspecifics as strong selective filters [22, 29]. Habitat effects were weaker but significant, suggesting that background and light environment modulate colour evolution in concert with behavioural and social factors. These results demonstrate that different ecological dimensions shape colour and morphology through partially distinct, yet interacting, selective pathways.

### Integrated phenotypes and adaptive landscapes

Despite distinct ecological drivers, colour and morphology exhibit significant evolutionary integration across pomacentrids. Six colour–shape axis pairs evolved in concert, forming partially integrated “trait syndromes” that link locomotion, feeding mechanics, and visual signalling [30]. For instance, body elongation covaried with anteroposterior pigmentation gradients, likely reflecting coordinated selection balancing crypsis, conspicuousness, and locomotor performance in heterogeneous reef light environments [3, 14].

Such covariation aligns with shared developmental constraints: pigment cell differentiation and craniofacial morphogenesis both derive from neural crest lineages, potentially coupling pigmentation and skeletal patterning through conserved regulatory networks [31, 32]. Quantitative genetic theory predicts that such integration channels evolution along predictable trajectories, producing recurrent syndromes under similar selective regimes [33, 34].

These integrated phenotypes imply a rugged adaptive landscape, where ecological opportunity interacts with developmental bias to constrain evolution along predictable trajectories. Supporting this view, multi-optimum OU models (including diversity dependence in IndoPacific clades) outperformed neutral alternatives, revealing repeated convergence on a finite set of viable phenotypic combinations shaped by both selection and constraint [35, 36].

### Predictability and repeatability in reef-fish evolution

The pervasive convergence observed across pomacentrid colour and morphology underscores the predictability of evolution in ecologically structured systems [37, 38]. While phylogenetic history constrains the limits of phenotypic variation, similar ecological contexts repeatedly draw unrelated lineages toward comparable adaptive peaks. Clownfishes and other anemone-associated damselfishes exemplify this determinism, repeatedly evolving distinctive colour syndromes and body forms in response to a shared symbiotic niche [39]. More broadly, the recurrence of similar ecomorphs across independent clades highlights the limited number of viable ecological strategies afforded by coral-reef environments [40, 41].

## Conclusion

By integrating continuous colour spaces, morphology, and ecology within a phylogenetic framework, we provide the most comprehensive macroevolutionary analysis of pomacentrid phenotypes to date. Our results show that damselfish diversification is best described not as a single radiation, but as a series of repeated adaptive episodes shaped by ecological opportunity, convergence, and partial trait integration. Colour and morphology evolve through distinct but coordinated pathways, producing recurrent phenotypic syndromes that reflect shared ecological pressures and constraints. These findings establish Pomacentridae as a powerful model for understanding how complex, integrated phenotypes evolve in structured environments and offer general insights into the predictability of evolution in adaptive radiations.

## Methods

### Time calibrated phylogeny

We utilized the latest well-resolved time-calibrated phylogenetic tree from McCord et al. 2021, including 345 out of the 422 extant pomacentrid species. Subsequently, the phylogenetic tree was pruned using the *ape* R package [42] by excluding *Chromis sanctaehelenae, Pomacentrus bangladeshius*, and *Pycnochromis bami* and retaining 343 species for which we had comprehensive colouration, morphological, and ecological data (Supplementary Figure 1).

### Colouration and morphological datasets

We compiled 1,183 lateral-view images of adult pomacentrids representing 343 species from multiple reef-fish image databases. For each species, one to five high-quality images were selected, excluding juveniles and any specimens with obstructed views, distortion, or poor colour fidelity. Image quality was assessed manually based on focus, resolution, and exposure. Minimal post-processing was applied to standardise appearance, including underwater colour correction using contrast-limited adaptive histogram equalization (CLAHE) in LAB colour space, adaptive histogram equalization for white balancing, saturation enhancement, and denoising, implemented in *OpenCV* [43] and *scikit-image* [44].

A total of 46 morphological landmarks were digitised on each image using FIJI [45] (Supplementary Figure S1). Landmark configurations were aligned using Procrustes superimposition in *geomorph* [46], producing a mean shape to which all specimens were registered. Missing pixel values generated during alignment were interpolated using surrounding pixel information.

Colouration was quantified using principal component analysis (PCA) on the aligned RGB images. Images were standardised for resolution and spatial extent, and the red, green, and blue channels were concatenated into a single vector per image. PCA was applied to the resulting matrix, and the first eight components were retained. Species-level scores were calculated by averaging image-level scores, yielding a single multivariate position for each species in colour space.

For morphology, fin-ray landmarks were excluded, retaining homologous landmarks describing body outlines, fin insertions, eye position, opercle, and pre-opercle. From the aligned landmarks, we derived ecologically relevant ratios (body, head, snout, eye-height, peduncle) and snout angle, reflecting functional variation in swimming performance, feeding, and vertical positioning [47, 48]. Species means were calculated, and phylogenetic PCA was performed on 11 traits using phyl.pca in *phytools* to account for shared ancestry. The first eight axes, explaining 92.4% of total morphometric variation, were retained for subsequent analyses.

### Ecological data

We analysed three categories of ecological traits: diet, habitat, and interspecific interactions. Dietary ecotypes were taken from McCord et al. 2021 and classified as ‘benthic’ herbivory, ‘pelagic’ planktivory, or ‘intermediate’ omnivory. Habitat type was defined as the primary adult habitat based on *FishBase* records accessed via *rFishBase* [50] and corroborated with additional literature (Supplementary Data). To facilitate trait reconstruction, habitats were grouped into five classes: ‘sea anemones’ (clownfishes and anemone-associated species of *Amblypomacentrus* and *Dascyllus*), ‘freshwater’, ‘non-reef’ (e.g. seagrass, rubble, sand), ‘rocky reef’, and ‘coral reef’. Symbiotic interactions were classified into three categories: ‘mutualistic’ (e.g. anemonefishes), ‘commensalistic’ (e.g. farming damselfishes; [51]), and ‘free-living’ species without persistent interspecific associations.

### Partitioning of phenotypic variation and disparity through time

To assess the contribution of species identity and ecological factors to phenotypic structure, we conducted permutational multivariate analyses of variance (PERMANOVA) using the *vegan* package [52]. Analyses were performed on the combined colour and morphology PC datasets, first testing species identity as a grouping factor and subsequently evaluating diet, habitat, and symbiosis as predictors of species-level phenotypic composition.

To examine the temporal dynamics of phenotypic diversification, we conducted disparity-through-time (DTT) analyses on colour and morphology PC scores using the dtt function in *geiger* [53]. For each axis, empirical disparity trajectories were compared with null expectations generated from 1,000 Brownian-motion simulations. Deviations were quantified using the morphological disparity index (MDI), with positive values indicating greater within-clade overlap (convergence) and negative values indicating early among-clade partitioning (divergence).

### Ecological predictors of colour and morphological evolution

To assess the influence of ecology on colour and morphological evolution, we reconstructed the evolutionary histories of discrete ecological traits (diet, habitat, and symbiosis) and ancestral biogeographic ranges (see Supplementary Material & Methods), and used these reconstructions to define ecological regimes for downstream phylogenetic comparative analyses.

We tested whether ecological regimes predict multivariate trait variation using penalized-likelihood multivariate GLS in *mvMORPH* (mvgls) [54]. For the joint dataset (8 colour and 8 morphology PCs), we fitted global models (BM, OU, Early Burst [EB], Pagel’s *λ*) and regime-dependent models (BMM: multiple Brownian rates; OUM: multiple OU optima) based on consensus stochastic character maps generated from 100 replicate mappings under the best-supported transition model. Model fit was compared using the Expected Information Criterion (EIC) with 100 parametric bootstrap bias corrections [55]. Significance of ecological predictors was assessed using PERMANOVA with *Pillai*’s trace and 999 permutations.

Complementarily, we fitted univariate phylogenetic generalized least squares (PGLS) models for each PC axis using *nlme* [56]. Alternative phylogenetic correlation structures (BM, Pagel’s *λ*, OU) and ecological predictor sets (diet, habitat, symbiosis, and their combinations) were compared using AICc via *MuMIn* [57]. The best-supported model per PC was retained, coefficients and confidence intervals extracted, and *p*-values adjusted for multiple testing using false discovery rate (FDR) correction.

### Trait evolutionary models of colour and morphology

We fitted and compared alternative models of trait evolution for each colour and morphology principal component (PC1–PC8) using the *geiger* [53], *OUwie* [36], and *RPANDA* [58] R packages. Models included Brownian motion (BM), white noise (WN), single-optimum Ornstein– Uhlenbeck (OU), multi-optimum OU with discrete ecological regimes (OUM; diet, habitat, symbiosis), diversity-dependent models (DDlin, DDexp), and a matching-competition model (MC). For OUM models, ecological regime histories were derived from ancestral state reconstructions under the best-supported transition models. Model support was assessed using AICc and Akaike weights. Relative support among competing models was quantified using Evidence Ratios (ERs), calculated as the ratio of Akaike weights between the best-supported model and alternatives. ERs were classified as < 3 (indistinguishable), 3–20 (positive), 20–150 (strong), and > 150 (very strong), with higher values indicating increasingly decisive evidence for the best model.

To evaluate potential geographic heterogeneity while accounting for computational constraints, we re-fitted models within major biogeographic regions (IO, IAA, CPO, EPO, AO) using regional species subsets. Regional analyses were exploratory due to uneven lineage representation, and confirmed no tree-splitting bias by testing associations between bestmodel support and phylogenetic/trait properties (phylogenetic diversity, tree imbalance, Blomberg’s *K* and trait variance).

### Detection of trait shifts and convergent evolution

To identify convergent shifts in phenotypic optima (colouration and morphology) without predefined regimes, we applied *ℓ*1ou [59], which detects shifts in OU models using a regularized, data-driven approach. Analyses were performed separately for colour and morphology PCs, with the number of shifts selected using the phylogenetically adjusted Bayesian information criterion (pBIC). Robustness of inferred regimes was assessed with 100 bootstrap replicates, and shifts recovered in ≥ 70% of replicates were considered well supported.

### Phylogenetic associations between colour and morphology

To test whether morphology predicts colour variation while accounting for shared ancestry, we conducted univariate and multivariate PGLS analyses. Univariate PGLS regressed each colour PC against each morphological PC using gls in *nlme* [56], comparing Brownian motion, Pagel’s *λ*, and OU correlation structures and retaining the best model by AICc. Regression coefficients, *R*^2^, and *p*-values were extracted, with false discovery rate (FDR) correction applied within each colour PC.

To assess trait integration across axes, we fitted multivariate PGLS models using mvgls in *mvMORPH* [54], treating colour PCs as a multivariate response and morphology PCs as predictors. Alternative phylogenetic structures (BM, *λ*, OU, EB) were compared using the Expected Information Criterion (EIC), and significance was evaluated by PERMANOVA with *Pillai*’s trace and 999 permutations. Colour was modelled as the response to reflect the hypothesis that body shape provides the structural context for colour-pattern expression.

## Supplementary information

## Acknowledgements

We thank Matías García for his help in landmark setting and his invaluable contribution. We also appreciate the valuable contribution of Lucy Fitzgerald, Anna Marcionetti, Diego A. Hartasánchez and Thibault Latrille through their useful comments on the manuscript.

## Declarations

### Data and code availability

Data and scripts with instructions to reproduce the analysis and figures can be found at https://github.com/agarciaEE/DamselfishTraitEvolution2025 ensuring transparency and enabling the reproducibility of our research findings. Access to the full raw data will be provided through Zenodo repository (DOI:10.5281/zenodo.17225926) upon acceptance of the manuscript.

### Author contribution

AGJ, TG and NS designed the study. AGJ conducted the research and analyses and wrote the manuscript. TG and NS contributed to the interpretation of the analyses, and the writing of the manuscript. All authors participated in reviewing, correcting, and approving the final version of the manuscript.

