## Supplementary for "Colour and shape evolution reflect ecological specialisation in Pomacentridae"

**Abstract**

Damselfishes (Pomacentridae) display remarkable diversity in colouration and body form, yet the processes shaping this phenotypic variation remain poorly resolved. Our study aimed to characterise the evolution of these traits, evaluate their associations with ecological factors, and identify convergent patterns linked to ecological specialisation. Using image-based quantification of colour patterns, geometric morphometrics, and phylogenetic comparative methods across 343 species, we show that pomacentrid phenotypes are organised around a small number of dominant axes describing brightness, hue, contrast, body elongation, and cranial morphology. Both colour and morphology exhibit early bursts of evolutionary disparity, followed by recurrent lineage-specific radiations and widespread convergence toward similar adaptive optima across the phylogeny. Dietary ecotypes emerged as the strongest predictor of morphological diversification, whereas symbiotic and social regimes exerted the strongest effects on colour evolution. Despite these distinct ecological correlates, several colour and shape axes form partially integrated trait syndromes that evolve in concert. The pervasive convergence of colour–shape syndromes underscores deterministic components of reef-fish evolution and positions Pomacentridae as a model for understanding integrated phenotypic evolution.

**Keywords:** Convergent evolution, Colouration, Morphology, Pomacentridae, Macroevolution, Reef fish, Ecological adaptation, Trait integration, Adaptive radiation, Phenotypic diversification

#### Supplementary Material & Methods

We conducted all analyses in R [v.4.3.2; 1], using *phytools* [2] and *ape* [3] for phylogenetic data manipulation. Visualizations were produced with *ggplot2* [4], and tables and reports were generated with *knitr* [5] and *kableExtra* [6].

##### Ancestral state reconstructions

To test the influence of ecology on the evolution of morphology and coloration in downstream analyses, we reconstructed ancestral states (ASR) for three discrete ecological traits, diet ecotype, habitat type, and symbiotic behavior, using a combination of explicit transition-rate models and stochastic character mapping. All analyses were conducted in `R` using the *phytools* [2], *corHMM* [7], and *geiger* [8] packages.

For each trait, we fitted a series of continuous-time Markov models that included standard configurations with equal rates (ER), all rates different (ARD), symmetric rates (SYM), and directionally constrained variants (DIR1–DIR3). In addition, we implemented biologically informed transition matrices ("CONSERVATIVE" and "RELAXED" models) tailored to each trait to capture plausible evolutionary constraints not represented in the standard models. For instance, in the *DietEcotype* and *Symbiosis* traits, direct transitions between extreme states (e.g., benthic ↔ pelagic, or free-living ↔ mutualistic) were initially forbidden under the conservative model and allowed as rare jumps under the relaxed model. For the *Habitat* trait, the transition matrix reflected the hierarchical and asymmetric nature of habitat evolution within Pomacentridae: stepwise transitions among marine habitats (non-reef ↔ rocky-reef ↔ coral-reef) were permitted, while direct transitions between freshwater and fully marine or symbiotic habitats were restricted. This directional constraint reflects the well-supported marine origin of damselfishes [9] and the very low probability of secondary colonisations of marine habitats from freshwater lineages [10].

Each model was fitted using `fitMk`, and the best-supported model was selected according to the corrected Akaike Information Criterion (AICc; 11). For selected models, we estimated both joint and marginal probabilities of ancestral states with `corHMM`, assuming a "FitzJohn" root prior. We further performed stochastic character mapping to quantify uncertainty in the reconstruction process. Using the best-fit transition matrix, we generated 1,000 stochastic maps per trait with `make.simmap` using fixed maximum-likelihood estimates.

The resulting ancestral reconstructions were subsequently used in phylogenetic comparative models (PCMs) to test evolutionary trait associations between ecological, morphological, and colour dimensions. Specifically, ASR-informed transition histories provided the discrete ecological regimes under which trait evolution models were fitted, enabling assessment of correlated trait evolution.

##### Ancestral biogeographic ranges

Ancestral biogeographic reconstructions of the Pomacentridae family were carried out using the R package *BioGeoBEARS* [12]. We classified species current distribution into five bioregions: Indian Ocean (IO), Indo-Australian Archipelago (IAA), Central Pacific (CPO), East Pacific (EPO) and Atlantic Ocean (AO). We performed a stratified analysis in which biogeographic reconstruction was divided and conducted in five time intervals corresponding to the following geological epochs: Eocene (56-34 Mya), Oligocene (34-23 Mya), Miocene (23-5 Mya), Pliocene (5-3 Mya) and Pleistocene-Holocene (3-0 Mya). A sixth region, Tethys Sea (TS) was allowed from the Eocene to the Oligocene, and then deactivated following the geological closure and disappearance of the Tethys sea. Similarly, IAA region was only activated during the Pliocene and Pleistocene-Holocene epochs because IO and CPO were widely connected before the Pliocene. For each time interval, we created a distance matrix containing distances among bioregions. Distances were estimated using geological coral reef habitat maps of each time period [13]. Pairwise geodesic distances between coral suitable areas of each bioregion were calculated using the function `geoDist` from the *oce* R package [14]. We conducted ancestral biogeographic reconstruction for pomacentrids using maximum likelihood estimation to determine parameters for DEC, DEC+J, DIVA, DIVA+J, BAYAREA, and BAYAREA+J models. We selected the optimal model based on AICc and LRT criteria. Subsequently, we employed Biogeographic Stochastic Mapping (BSM) with the chosen model, generating 50 stochastic maps illustrating the co-occurrence between lineages across the phylogenetic tree.

#### Supplementary Results

##### Ancestral reconstructions of ecological traits and biogeographic ranges

The best-fitting Mk models for the three ecological traits showed distinct and asymmetric transition structures (Supplementary Figure 10). For diet, the CONSERVATIVE model yielded the highest likelihood (logL = –216.95; AICc = 442.02). Transitions were most frequent from the intermediate state toward pelagic ($q_{I \to P} = 2.17$) and benthic ($q_{I \to B} = 1.45$) ecotypes, while direct benthic–pelagic transitions were disallowed. Stationary frequencies were $\pi_B = 0.10$, $\pi_I = 0.85$, and $\pi_P = 0.05$. For habitat, the RELAXED model was best supported (logL = –339.88; AICc = 698.29), with high rates from non-reef to rocky-reef ($q_{NR \to RR} = 2.33$) and coral-reef ($q_{NR \to CR} = 2.01$), and from rocky-reef to coral-reef ($q_{RR \to CR} = 1.95$). Transitions from freshwater or sea-anemone habitats were absent. Stationary frequencies were $\pi_{NR} = 0.23$, $\pi_{RR} = 0.58$, and $\pi_{CR} = 0.19$. For symbiosis, the RELAXED model provided the best fit (logL = –108.86; AICc = $227.90$). The highest transition rate occurred from commensalistic to free-living ($q_{C \to F} = 1.33$), with lower rates from free-living to commensalistic ($q_{F \to C} = 0.26$) or mutualistic ($q_{F \to M} = 0.06$). Stationary frequencies were $\pi_F = 0.42$, $\pi_C = 0.58$, and $\pi_M \approx 0.00$.

Ancestral reconstructions based on these best-fitting models (Supplementary Figure 11) inferred an intermediate diet for the pomacentrid ancestor, with frequent intermediate $\leftrightarrow$ pelagic and intermediate $\to$ benthic transitions (Supplementary Figure 12). Rocky-reef habitats were estimated as the most probable ancestral state, with dominant rocky-reef $\to$ coral-reef and coral-reef $\to$ non-reef transitions (Supplementary Figure 13). Commensalistic symbiosis was inferred as the ancestral state of pomacentrids, with balanced transitions between commensalistic and free-living conditions. Additional independent transitions toward mutualism occurred in the clownfish clade and in the sister pairs *Amblypomacentrus breviceps*–*A. clarus* and *Dascyllus trimaculatus*–*D. albisella* (Supplementary Figure 14).

For the biogeographic reconstruction, we selected the "DEC+J" model —the best-fitting model based on AICc (logL = -573.882; AICc = 1153.850; weight = 0.999)— to infer ancestral range evolution in pomacentrids. The results indicated a widespread origin of the clade, occurring in the Tethys Sea and the Central and Eastern Pacific Ocean followed by a rapid cladogenetic dispersal towards the Indian Ocean (Supplementary Figure 15). Anagenetic dispersal events explained most of the biogeographic events in pomacentrids, followed by narrow sympatry (or parapatry). Estimates of founder, vicariance and nested sympatry events were relatively low, with an average of 30, 30 and 70 counted events, respectively, over the 50 stochastic reconstructions (Supplementary Figure 16).

##### Patterns of phenotypic variance and phylogenetic dependence

Species identity explained a large proportion of variation in colour patterns ($R^2 = 0.703$, $F = 5.54$, $p = 0.001$), and similarly accounted for substantial differences in morphometric traits ($R^2 = 0.597$, $F = 3.588$, $p = 0.001$). For colour traits, habitat accounted for the largest share ($R^2 = 0.021$, $p = 0.005$), followed by diet ($R^2 = 0.013$, $p = 0.005$) and symbiosis ($R^2 = 0.007$, $p = 0.025$). For morphology, ecological predictors explained slightly more variance overall, with diet emerging as the strongest factor ($R^2 = 0.047$, $p = 0.001$), followed by habitat ($R^2 = 0.029$, $p = 0.001$), while symbiosis showed no detectable effect ($R^2 = 0.002$, $p = 0.493$).

#### Comparative analyses of colour and morphology under ecological regimes

##### *Colour variation.*

Univariate phylogenetic regressions revealed limited but significant ecological effects on colour traits (Supplementary Table 7). Most colour PCs were best explained by null models, indicating weak large-scale ecological structure. However, PC4 (dorsoventral contrast) showed strong associations with both habitat and diet under a *Habitat+Diet* model with Pagel's $\lambda$ ($\lambda \approx 0.00$). Species inhabiting sea-anemone habitats ($\hat{\beta} = 0.55$, $t = 3.07$, $p_{FDR} = 0.007$) and benthic feeders ($\hat{\beta} = 0.48$, $t = 6.77$, $p_{FDR} < 0.001$) displayed higher PC4 scores, suggesting enhanced dorsoventral pigmentation contrasts in these ecological contexts. PC5 (vertical colour asymmetry) was best fit by an OU model ($\alpha = 1.04$), though no ecological predictors were significant.

Symbiotic lifestyle was the best-supported predictor of colour multivariate variation. The symbiosis model under Pagel's $\lambda$ yielded the lowest EIC (31,043.720; weight = 1.00) and was significant in permutation MANOVA ($Pillai = 0.112, p = 0.005$; Supplementary Table 11). The fitted model indicated a moderate phylogenetic signal ($\lambda = 0.48$) and large, axis-specific differences among symbiotic categories.

Relative to free-living species, commensalistic taxa exhibited lighter colour tones (PC1: +28.66), reddish and rostral bar intensity (PC2: +53.63), core-body shading (PC3: -23.43), increased antero-dorsal pigmentation (PC4: –18.06, and PC5: –23.39), opposite segmentation patterns (PC7: +14.99), and brighter peduncle pigmentation (PC8: +4.27). Mutualistic taxa exhibited stronger directional shifts, including a large positive effect on PC2 (+97.83), indicating strong reddish coloration and rostral white bar presence; and strong negative effects on PC3 (–43.43), PC4 (–67.01) and PC6 (–42.27), associated respectively with core-body shading, increased dorsal pigmentation, and rostral markings. Additional moderate shifts occurred on PC1 (–11.00; overall brightness), PC5 (+19.77; caudal pigmentation), PC7 (–11.16; barred segmentation) and PC8 (–21.30; ventral shadings). Diet and habitat also produced significant MANOVA effects ($p < 0.05$) but had substantially weaker support ($\Delta EIC > 12$), and the second-ranking overall model was the $\lambda$-structured null model ($\Delta EIC = 19.7$), indicating considerable phylogenetic structure in colour variation.

##### *Morphological variation.*

Univariate PGLS analyses revealed weak ecological structuring of body-shape variation (Supplementary Table 8). Among all axes, only PC5 —primarily associated with pelvic fin and snout proportions— showed significant support for a *DietEcotype* model under a Pagel's $\lambda$ framework ($\lambda = 0.115$). Pelagic feeders exhibited higher PC5 scores ($\hat{\beta} = 0.415$, $t = 2.82$, $p_{FDR} = 0.010$), indicating subtle morphological differentiation linked to feeding ecology. All remaining PCs were best explained by null models, consistent with largely neutral or weakly structured morphological diversification across ecological gradients.

Multivariate morphological variation was best explained by diet. The diet model with Pagel's $\lambda$ achieved the strongest fit (EIC = 22,159.287; weight = 1.00) with a moderate phylogenetic signal ($\lambda = 0.58$) and was supported by permutation MANOVA ($Pillai = 0.177, p = 0.001$; Supplementary Table 10). Intermediate-diet species differed from the benthic diet category along several axes, with positive effects on PC1 (+2.79; more elongated bodies with taller peduncles and elevated eyes) and PC2 (+2.15; larger eyes and increased snout angle), and negative effects on PC3 (–2.95; deeper heads with larger eyes), PC4 (–2.82; steeper snout inclination and more anterior pelvic-fin placement), PC6 (–3.52; longer anal fins and more streamlined profiles) and PC7 (–1.07; more streamlined head shapes with reduced head ratio). Pelagic taxa showed larger deviations, including strong positive displacement on PC1 (+6.18; markedly elongated bodies with narrower peduncles) and strong negative effects on PC3 (–5.65; deeper heads with larger eyes) and PC6 (–4.69; elongated anal fins and strongly compressed body shapes), alongside additional smaller effects across PC2 (+1.58; larger eyes and steeper snout angle), PC4 (–1.63; steeper snout inclination), PC5 (+1.58; larger pelvic fins), PC7 (–1.30; streamlined head profiles), PC8 (+0.14; elevated eye position and increased body height) and PC9 (–0.65; high-set eyes with shorter snouts). Symbiosis and habitat also yielded significant MANOVA results but were markedly less supported ($\Delta EIC = 21.1$ and $50.0$, respectively). Regime-dependent OUM models improved over BM and BMM fits but did not outperform ecological predictors under Pagel's $\lambda$.

#### Model comparison of colour and morphological trait evolution

Regional analyses revealed heterogeneity in model support across biogegraphic regions (Supplementary Figure 6b and 8). For both colour and morphology, OUM models predominated across all regions, while density-dependent processes were recovered in the Indo-Australian Archipelago (IAA) and Central Pacific Ocean (CPO) for several colour and morphological axes. WN models were often best supported in the Eastern Pacific Ocean (EPO) and particularly the Atlantic Ocean (AO) for most of colour and morphological axes, indicating a lack of detectable phylogenetic signal. Analyses of structural bias confirmed that these outcomes were not artefacts of tree-splitting or uneven trait sampling across regions, as neither phylogenetic diversity, tree imbalance, nor trait variance systematically biased model support (Supplementary Figure 9).

### Supplementary Discussion

#### Historical and adaptive foundations of pomacentrid diversification

Ancestral reconstructions and biogeographic models reveal that the early evolution of Pomacentridae was deeply shaped by shifting palaeoenvironmental contexts. The ancestral pomacentrid likely inhabited rocky-reef environments, displayed intermediate feeding strategies, and engaged in commensal interactions —a ecological scenario compatible with the Tethyan and Indo-Pacific origins of reef-fish faunas [9, 13, 15, 16]. From this generalist ancestor, recurrent transitions toward coral-reef specialisation and pelagic or benthic diets accompanied the Miocene expansion of coral habitats, catalysing major shifts in body form and colouration [17–20]. However, the asymmetry and rarity between certain transitions (e.g. benthic–pelagic) suggest the presence of ecological or developmental "barriers" between these modes, a pattern previously observed in jaw functional evolution [21] and more broadly across water-column transitions in reef fishes [22]. Frequent intermediate or omnivorous states, however, appear to have acted as gateways facilitating diversification between extremes, explaining why similar morphotypes and colour syndromes recur across distantly related lineages that occupy transitional or mixed habitats, such as lagoonal reefs and outer-reef slopes.

Ancestral-state estimates of symbiosis indicate a moderately elevated probability of commensalism at the family root, possibly reflecting the widespread and repeated evolution of commensalism across independent lineages that pulls the likelihood of this state deeper in the tree. Yet behavioural observations support the plausibility of early proto-symbiotic behaviours —such as algal farming or the opportunistic use of shelter-providing organisms— that precede coral-reef colonisation [23, 24]. These versatile strategies, combined with an omnivorous diet, may have conferred ecological flexibility that enabled subsequent specialisation and ultimately the evolution of obligate anemone mutualism in clownfishes [25, 26].

Biogeographic reconstructions under the `DEC+J` model point to an early widespread distribution across the Tethys and Central Pacific, followed by dispersal into the Indian Ocean and diversification as the Indo-Australian Archipelago emerged [13, 16, 27]. Fossil evidence from the Eocene Monte Bolca sites confirms damselfish presence in Tethyan reefs [16], supporting a scenario in which early pomacentrids occupied broad marine habitats before expanding into modern coral reefs. Similar origins in other lineages [28, 29] suggest that coral-reef colonisation was a secondary phase driven by increasing reef complexity during the Eocene–Oligocene [30, 31]. The combination of widespread ancestral ranges and high dispersal capacity likely promoted parallel adaptation to similar environmental gradients, reinforcing the recurrent evolution of comparable ecological strategies across lineages. Later contractions and fragmentation of reef habitats, coupled with vicariance and founder events, further amplified regional endemism and contributed to the exceptional species richness of pomacentrids within the Indo-Pacific hotspot [9, 27, 28].

#### Interpretational scope and future extensions

While our analyses capture broad macroevolutionary patterns in pomacentrid phenotypes, several sources of ecological, developmental, and biogeographic complexity necessarily fall outside the present framework.

##### Limits of categorical ecological classifications

Our ecological reconstructions necessarily rely on discrete trait categories for diet, habitat, and symbiosis, which simplify a continuum of ecological strategies. Pomacentrids exhibit fine-scale variation in microhabitat use, ontogenetic diet shifts, and behavioural plasticity that are not fully captured by categorical coding [32, 33]. For example, several nominally benthic or intermediate feeders exhibit seasonal or context-dependent foraging strategies, while habitat associations may vary across life stages or geographic regions [34, 35]. Such ecological complexity likely contributes to the substantial residual variance observed across colour and morphological axes and may underlie lineage-specific deviations from inferred adaptive optima.

Future work integrating continuous environmental variables (e.g. depth, light spectra, habitat complexity), behavioural traits, and ontogenetic data could refine ecological regime definitions and improve resolution of eco-phenotypic associations [36, 37]. This would allow more explicit testing of how fine-scale ecological gradients interact with colour and morphological evolution beyond the limits of categorical predictors.

##### Ontogeny, sexual dimorphism, and plasticity in colour evolution

Our analyses focus on adult phenotypes and therefore do not explicitly account for ontogenetic colour change, sexual dimorphism, or context-dependent plasticity. Several pomacentrid lineages exhibit pronounced juvenile–adult colour transitions or sex-specific patterns that serve distinct ecological or social functions, including predator avoidance, dominance signalling, or mate recognition [38, 39]. Such dynamics may generate transient or labile colour states that are difficult to capture in macroevolutionary frameworks based on adult phenotypes alone.

Incorporating ontogenetic trajectories or sex-specific colouration into comparative models could reveal additional axes of variation and help disentangle selection acting at different life stages [40]. These approaches would also clarify whether convergent adult colour patterns arise through shared developmental pathways or through distinct ontogenetic routes.

#### Developmental and genetic bases of trait integration

The partial integration observed between colour and morphology suggests shared developmental or genetic constraints linking pigmentation and craniofacial traits. In teleosts, pigment cell differentiation, migration, and patterning are closely associated with neural crest development, which also contributes to craniofacial morphology [41, 42]. This raises the possibility that correlated trait evolution reflects pleiotropy, shared regulatory pathways, or developmental coupling.

While our analyses cannot resolve the mechanistic basis of this integration, emerging genomic resources for pomacentrids and other reef fishes offer opportunities to test whether repeated colour–shape syndromes arise through conserved molecular architectures or distinct genetic solutions [e.g., 43]. Comparative genomic or transcriptomic analyses could reveal whether phenotypic convergence masks deeper developmental divergence.

#### Geographic structure and biogeographic contingencies

Although our phylogenetic framework accounts for shared ancestry, it does not explicitly model geographic structure or historical connectivity among reef systems. Biogeographic history may strongly influence the availability of ecological opportunities and the tempo of adaptive diversification. Indo-Pacific hotspots are characterised by high species richness and ecological saturation, whereas peripheral regions such as the Atlantic show lower diversity and more recent radiations following isolation events [44, 45].

Explicitly integrating geographic range evolution with trait dynamics could clarify how regional context shapes adaptive landscapes and whether convergence differs systematically across biogeographic provinces. Our analyses provide an exploratory indication of such effects, as diversity-dependent signals were more frequently detected in species-rich Indo-Pacific clades, consistent with ecological saturation modulating evolutionary rates [46, 47].

Support for diversity-dependent models on a subset of trait axes suggests that interspecific interactions and niche packing may influence evolutionary dynamics. However, diversity dependence can arise from multiple processes, including competition, environmental change, or methodological artefacts [48]. These signals should therefore be interpreted cautiously and evaluated alongside independent ecological and biogeographic evidence. Future work integrating fossil data, explicit competition models, or community-level trait distributions could help disentangle these mechanisms.

#### Generality beyond Pomacentridae

While Pomacentridae represent an ideal system for studying integrated phenotypic evolution, the extent to which these patterns generalise across reef fishes remains an open question. Other clades with different life histories, dispersal capacities, or sensory ecologies may exhibit weaker trait integration or alternative adaptive trajectories [49, 50].

Applying comparable multivariate and phylogenetic frameworks across reef-fish families will be essential to determine whether convergence toward integrated colour–morphology syndromes reflects a general property of reef-fish evolution or a distinctive feature of pomacentrids. Such comparative extensions will help clarify the broader evolutionary rules governing phenotypic diversification in complex marine ecosystems.

#### Supplementary Figures

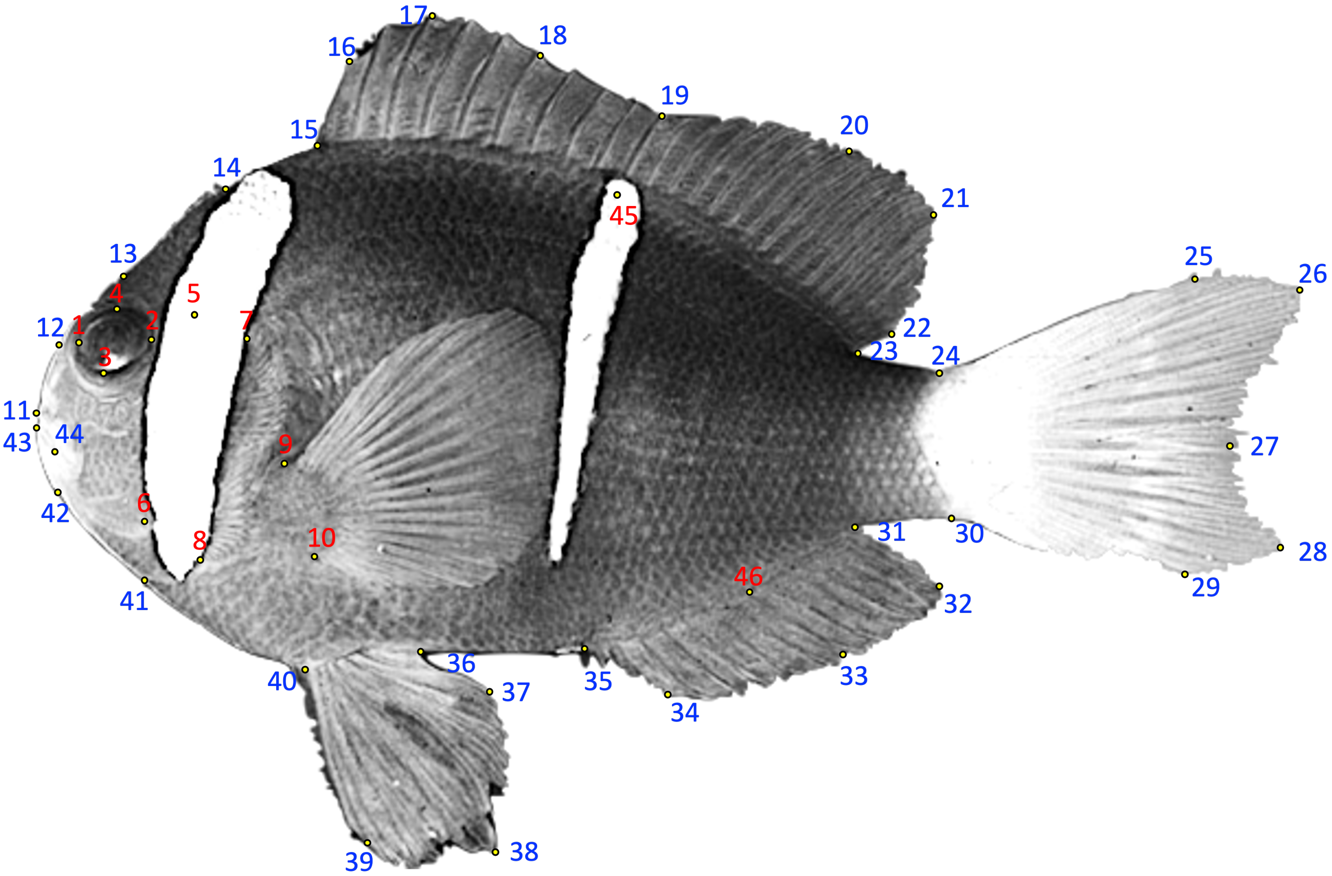


**Fig. 1**: Landmark-based geometric morphometric mapping of a damselfish specimen. The image shows a grayscale lateral view of a damselfish with manually placed anatomical landmarks used for geometric morphometric analysis. Landmarks are categorized into two sets: (1) Blue-numbered landmarks (1–44) represent external body shape landmarks capturing overall morphology, including head, fin, and body contours. (2) Red-numbered landmarks (1–10, 45–46) denote additional key anatomical reference points, specifically marking fin insertions, eye position, and stripe pattern boundaries. Small yellow dots indicate the exact placement of each landmark. This configuration facilitates quantitative shape analysis and comparisons across specimens or species.

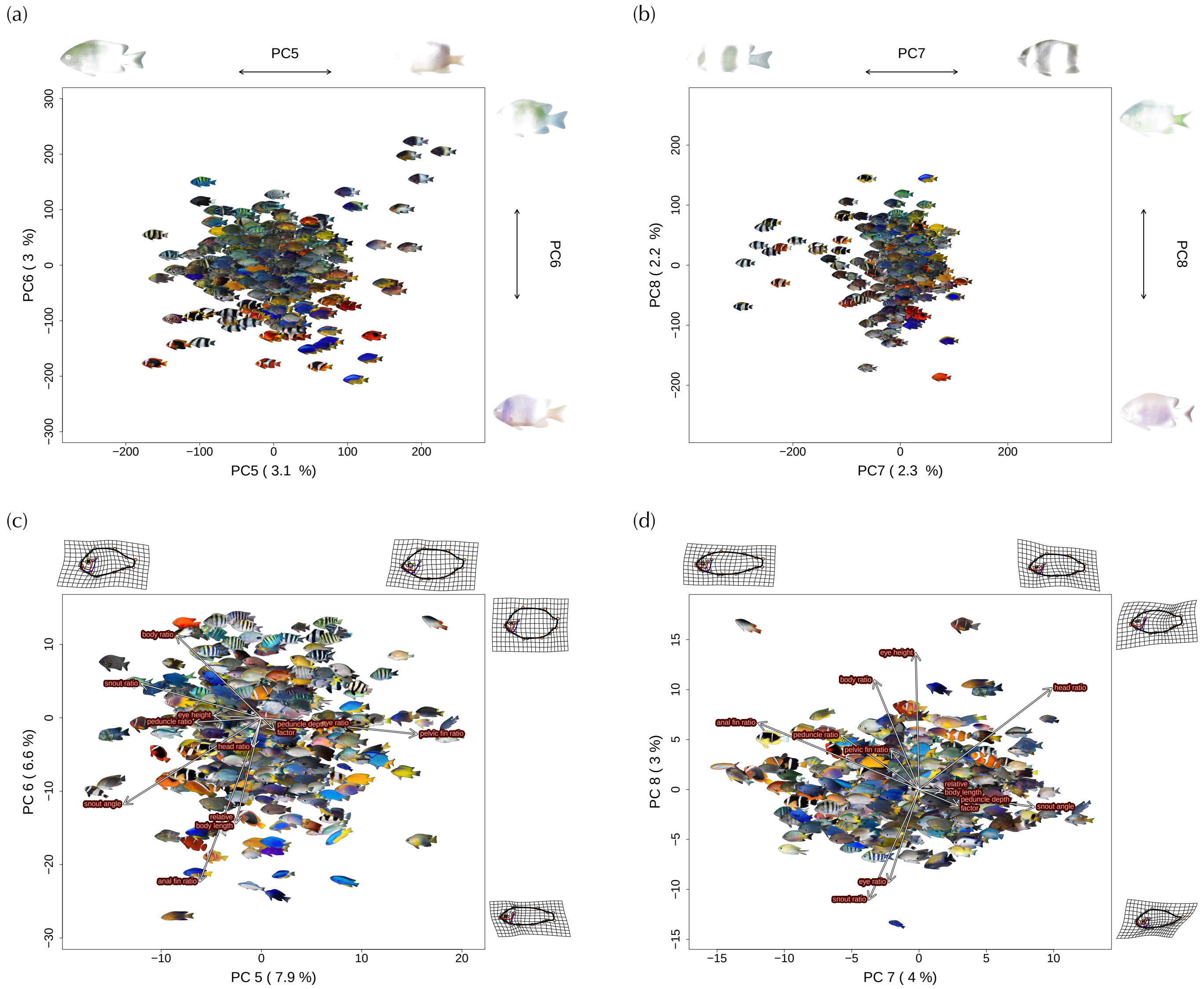


**Fig. 2**: Combined colour-pattern and body-shape principal component analyses (PC5–PC8) across 343 pomacentrid species. Panels (a) and (b) show colour PC5–6 (a) and colour PC7–8 (b), while panels (c) and (d) show the corresponding morpho PC5–6 (c) and morpho PC7–8 (d). For the colour PCA, each point represents species-averaged pixel coordinates, with reconstructed average images projected onto the morphospace. Reconstructed images on the plot margins display extreme colour pattern values along each PC axis. Pixel opacity scales with contribution to axis direction (note: white indicates zero contribution and may be indistinguishable from naturally white pattern elements). Variance explained for each axis appears on the axis labels. For the morphology PCA, raw images (background removed) are plotted instead of points. White arrows indicate eigenvector loadings of each morphometric trait, with orange labels at arrow tips. Variance explained for each axis appears on the axis labels. Marginal deformation grids show minimum and maximum shape changes along each PC, with orange landmarks and black outlines indicating major anatomical structures (body, eye, preopercle, and opercle contours).

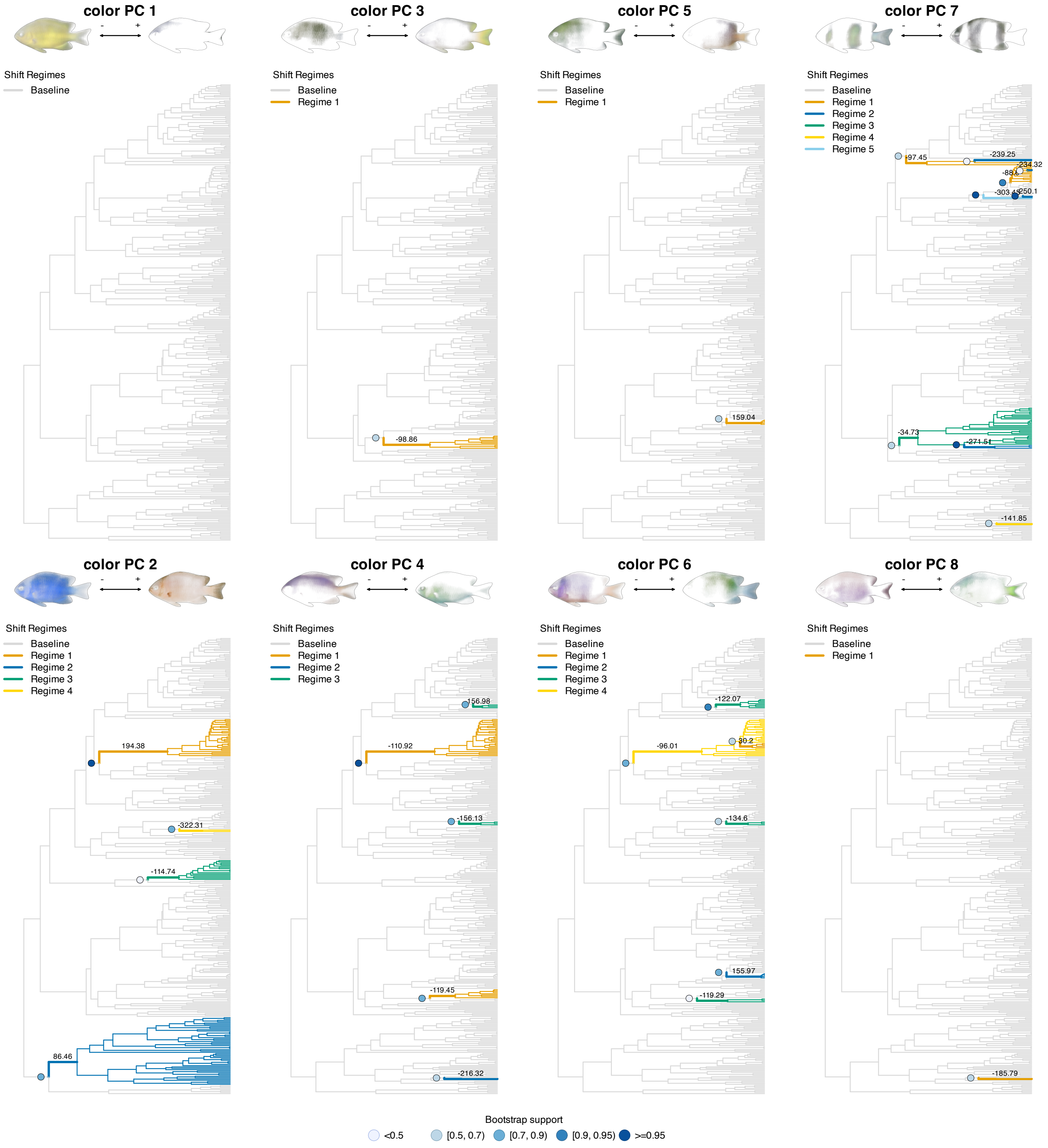


**Fig. 3**: $\ell$1ou-inferred adaptive regime shifts across the damselfish phylogeny based on univariate analyses of color principal component (PC) axes. At the top of each PC panel, reconstructed colour patterns illustrating the trait variation encoded by the axis, indicating the positive (+) and negative (–) extremes. Grey branches indicate the baseline (ancestral) regime, and coloured branches represent inferred shifts to alternative selective regimes. Identical colours across phylogenetically distant clades indicate convergent evolution toward similar trait optima. The estimated optimum value from $\ell$1ou is indicated at the start of each regime shift. Each panel includes a regime legend in the top-left corner, summarizing the number and identity of inferred regimes. Bootstrap support is shown at the start of each detected shift as a small filled circle whose colour indicates the categorical bin in the legend at the bottom (values denote the proportion of bootstrap replicates recovering the same shift). Shifts were detected based on the most conservative criterion 'pBIC'.

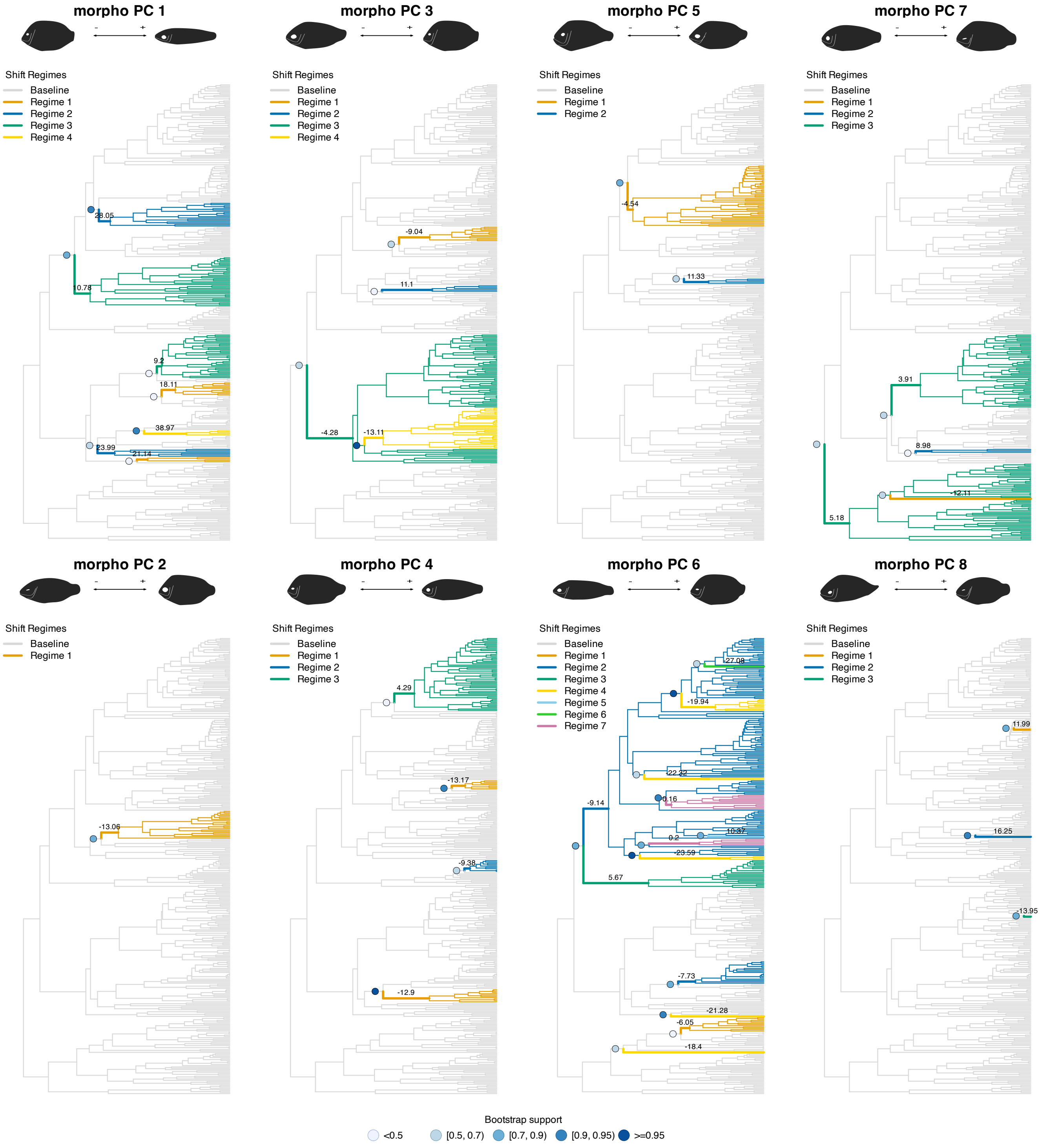


**Fig. 4**: `ℓ1ou`-inferred adaptive regime shifts across the damselfish phylogeny based on univariate analyses of morphological principal component (PC) axes. At the top of each PC panel, reconstructed morphological patterns illustrating the trait variation encoded by the axis, indicating the positive (+) and negative (–) extremes. Grey branches indicate the baseline (ancestral) regime, and coloured branches represent inferred shifts to alternative selective regimes. Identical colours across phylogenetically distant clades indicate convergent evolution toward similar trait optima. The estimated optimum value from `ℓ1ou` is indicated at the start of each regime shift. Each panel includes a regime legend in the top-left corner, summarizing the number and identity of inferred regimes. Bootstrap support is shown at the start of each detected shift as a small filled circle whose colour indicates the categorical bin in the legend at the bottom (values denote the proportion of bootstrap replicates recovering the same shift). Shifts were detected based on the most conservative criterion 'pBIC'.

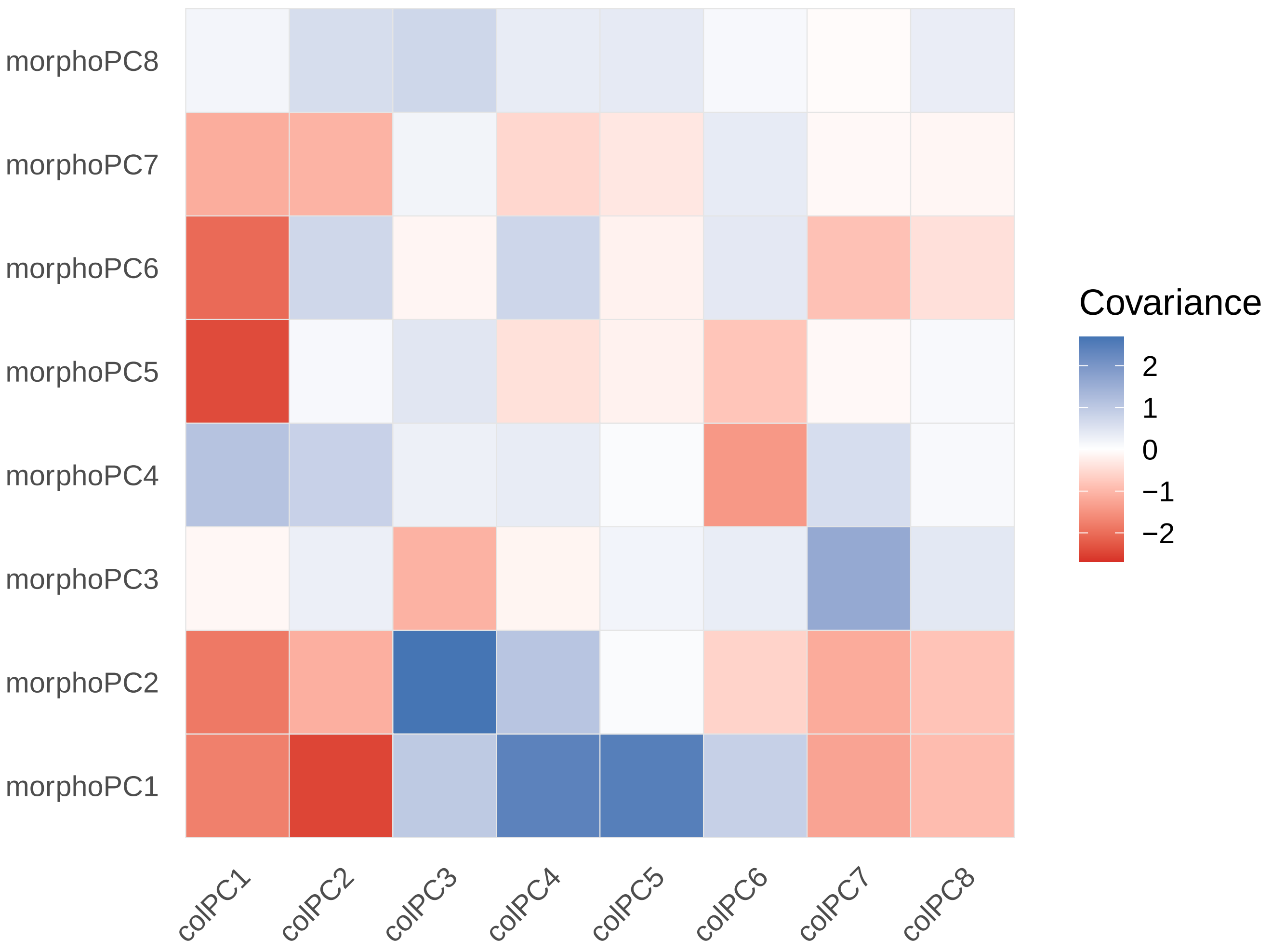


**Fig. 5**: Heatmap of pairwise covariances between colour and morphology principal components (PCs) estimated from the best–supported multivariate phylogenetic model (*mvGLS* with Pagel's $\lambda$, DietEcotype as predictor). Each cell displays the residual covariance between a colour PC (columns) and a morphology PC (rows) extracted from the model's trait covariance matrix. Positive values (blue coloured) indicate that colour and morphological traits tend to vary in the same direction across species, whereas negative values (red coloured) reflect opposing changes.

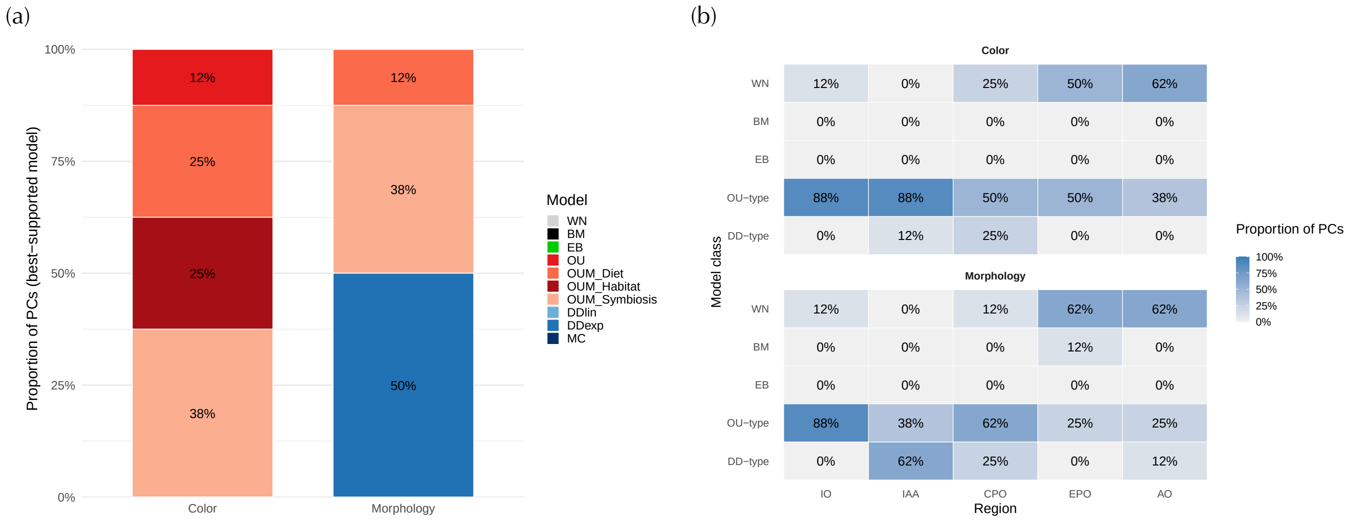


| Color | IO | IAA | CPO | EPO | AO |
|---|---|---|---|---|---|
| WN | 12% | 0% | 25% | 50% | 62% |
| BM | 0% | 0% | 0% | 0% | 0% |
| EB | 0% | 0% | 0% | 0% | 0% |
| OU−type | 88% | 88% | 50% | 50% | 38% |
| DD−type | 0% | 12% | 25% | 0% | 0% |

| Morphology | IO | IAA | CPO | EPO | AO |
|---|---|---|---|---|---|
| WN | 12% | 0% | 12% | 62% | 62% |
| BM | 0% | 0% | 0% | 12% | 0% |
| EB | 0% | 0% | 0% | 0% | 0% |
| OU−type | 88% | 38% | 62% | 25% | 25% |
| DD−type | 0% | 62% | 25% | 0% | 12% |

**Fig. 6**: Model support for trait evolution in Pomacentridae. (a) Overall model support across all principal components of colour and morphology traits. Bars show the proportion of principal component axes best supported by each evolutionary model (WN = white noise, BM = Brownian motion, OU-type = single- or multi-optimum Ornstein–Uhlenbeck models [OU, OUM by Diet, Habitat, or Symbiosis], and DD-type = diversity-dependent models [DDlin, DDexp, MC]). (b) Regional patterns of best-supported models. Heatmaps show the proportion of PCs within each region (IO = Indian Ocean, IAA = Indo-Australian Archipelago, CPO = Central Pacific Ocean, EPO = Eastern Pacific Ocean, AO = Atlantic Ocean) best explained by each model class. Grouped model families are indicated as in (a).

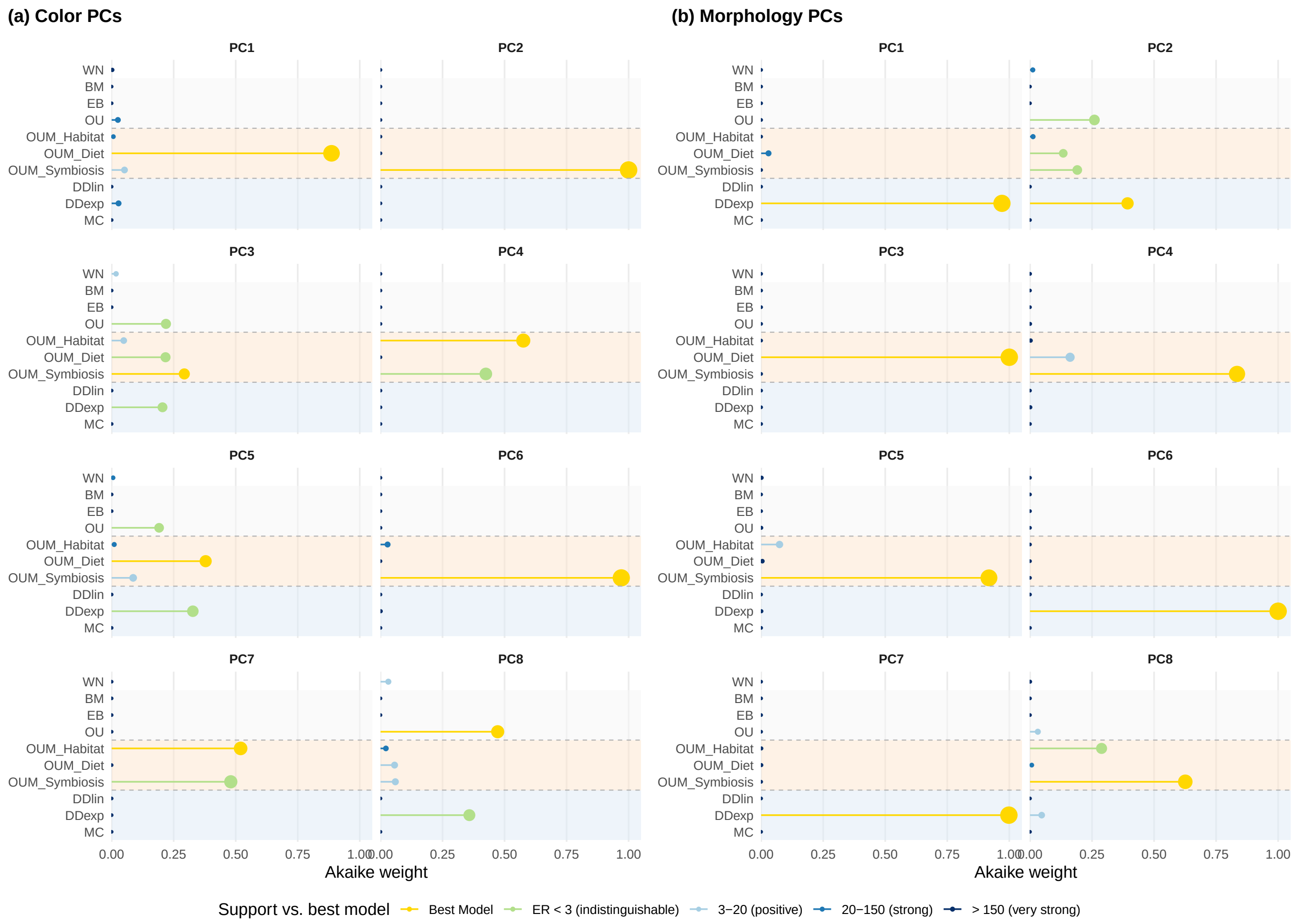


**Fig. 7**: Global model support across principal components (PC1–PC8) of colour (a) and morphology (b). Lollipop plots show the Akaike weights different evolutionary models fitted for each PC. Lollipop colors indicate relative evidence ratios (ERs) of alternative evolutionary models compared to the best-fitting model support grouped in categories: gold = best model; light green = ER < 3 (indistinguishable); light blue = 3–20 (positive); medium blue = 20–150 (strong); and dark blue = > 150 (very strong). Background shading groups model families: light grey = simple models (WN, BM, OU); light orange = multi-optima OUM models; and light blue = density-dependent (DD) models.

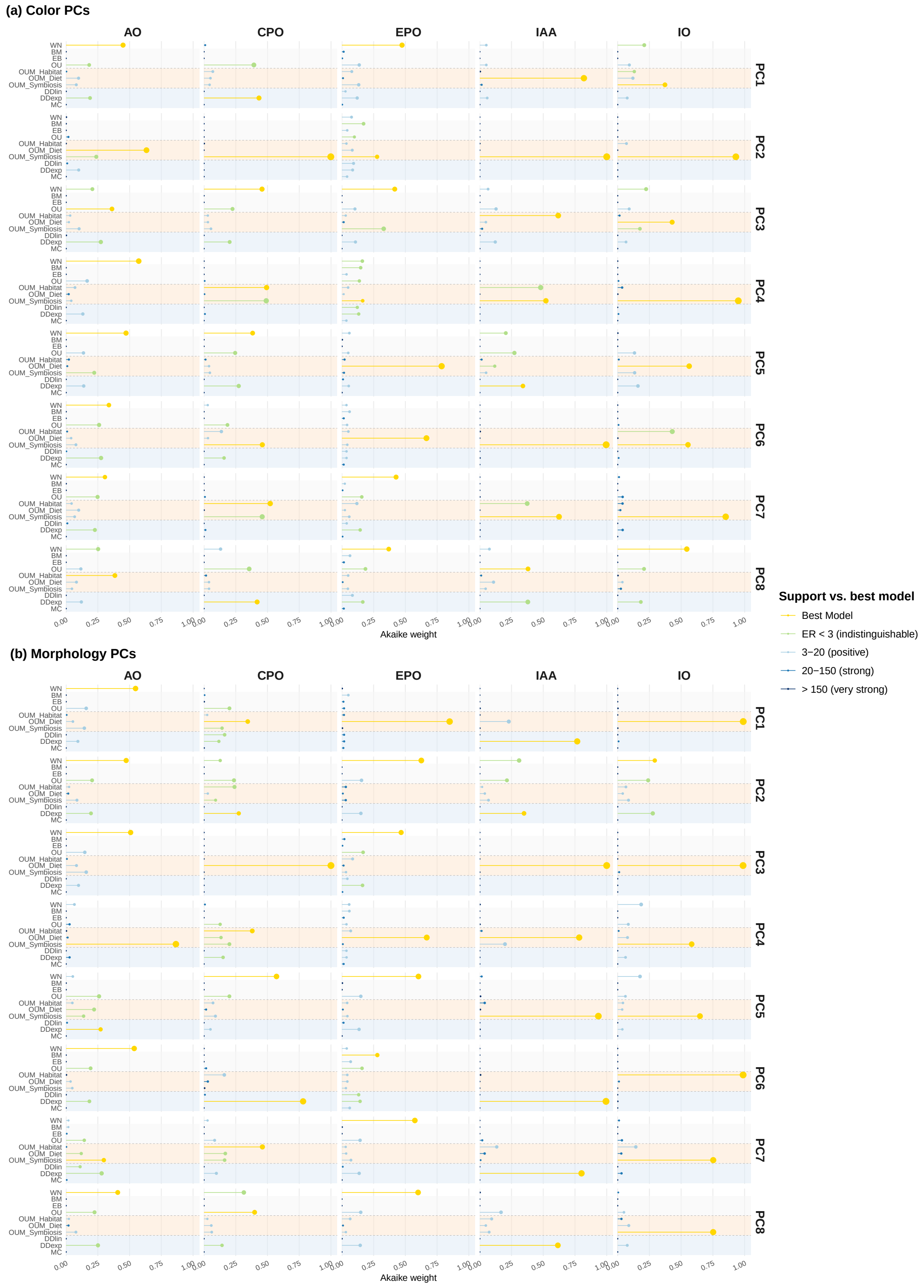


**Fig. 8**: Regional model support across biogeographic regions (AO = Atlantic Ocean, CPO = Central Pacific Ocean, EPO = Eastern Pacific Ocean, IAA = Indo-Australian Archipelago, IO = Indian Ocean). Panels show the Akaike weights of different evolutionary models fitted to colour PCs (a) and morphology PCs (b) across regions. Lollipop colors relative evidence ratios (ERs) of alternative evolutionary models compared to the best-fitting model grouped in categories: gold = best model; light green = ER < 3 (indistinguishable); light blue = 3–20 (positive); medium blue = 20–150 (strong); and dark blue = > 150 (very strong). Background shading groups model families: simple models (WN, BM, OU) in light grey, multi-optima OUM models in light orange, and density-dependent (DD) models in light blue.

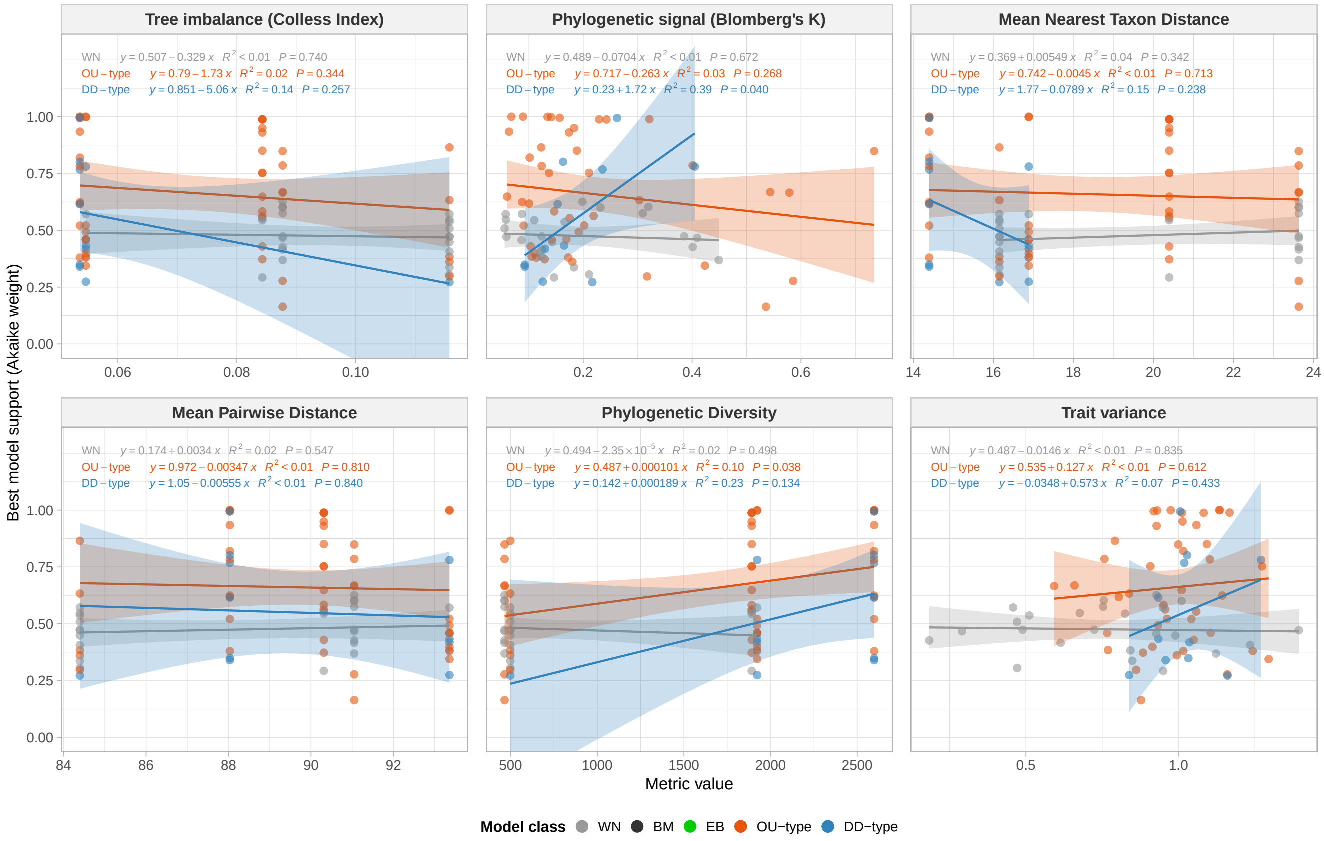


**Fig. 9**: Structural bias analysis of regional model support for colour and morphology univariate evolutionary model analyses. Each panel shows the relationship between the Akaike weight of the best-supported model (y-axis) and a tree- or trait-based metric (x-axis): tree imbalance, phylogenetic signal (Blomberg's $K$), mean nearest taxon distance (MNTD), mean pairwise distance (MPD), phylogenetic diversity (PD), and trait variance. All best-supported model weights across colour and morphology PC axes are included. Independent linear fits are shown for each model family (WN, BM, OU-type, and DD-type), with fitted equations, $R^2$, and $P$-values displayed at the top of each panel. Regression fits for BM model weights are not shown, as this model family was best supported in only one case across all colour and morphology PC axes. No consistent linear trends are detected, although weak positive associations are observed between Blomberg's $K$ and DD-type models, and between phylogenetic diversity and OU-type models, suggesting minor structural influences on regional model support.

#### (a) Diet ecotype

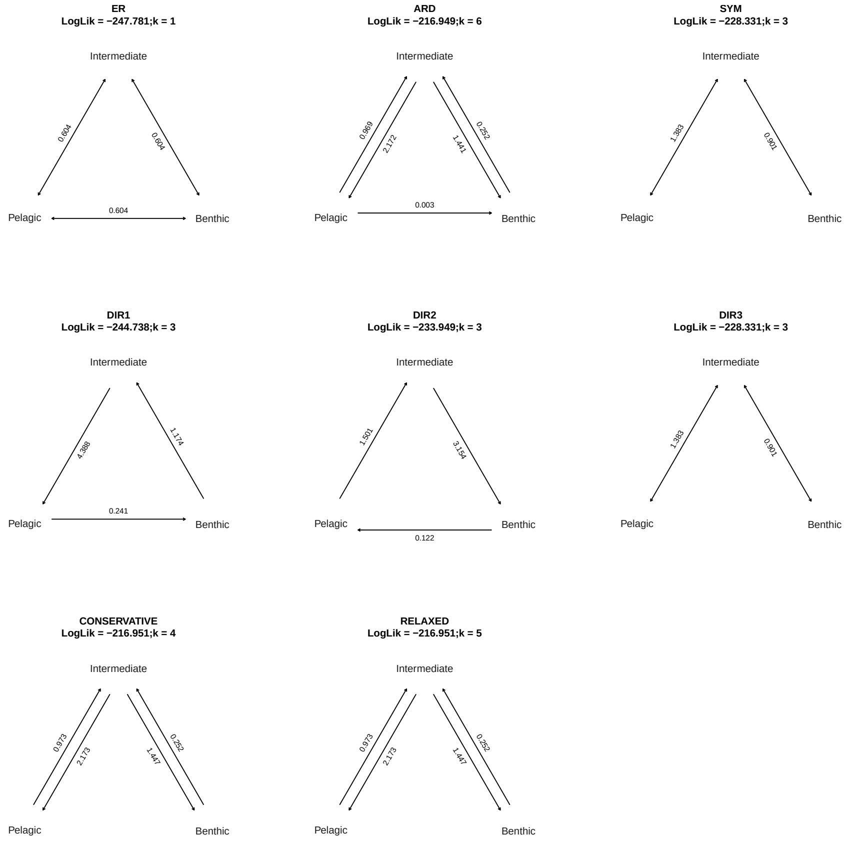


#### (b) Habitat type

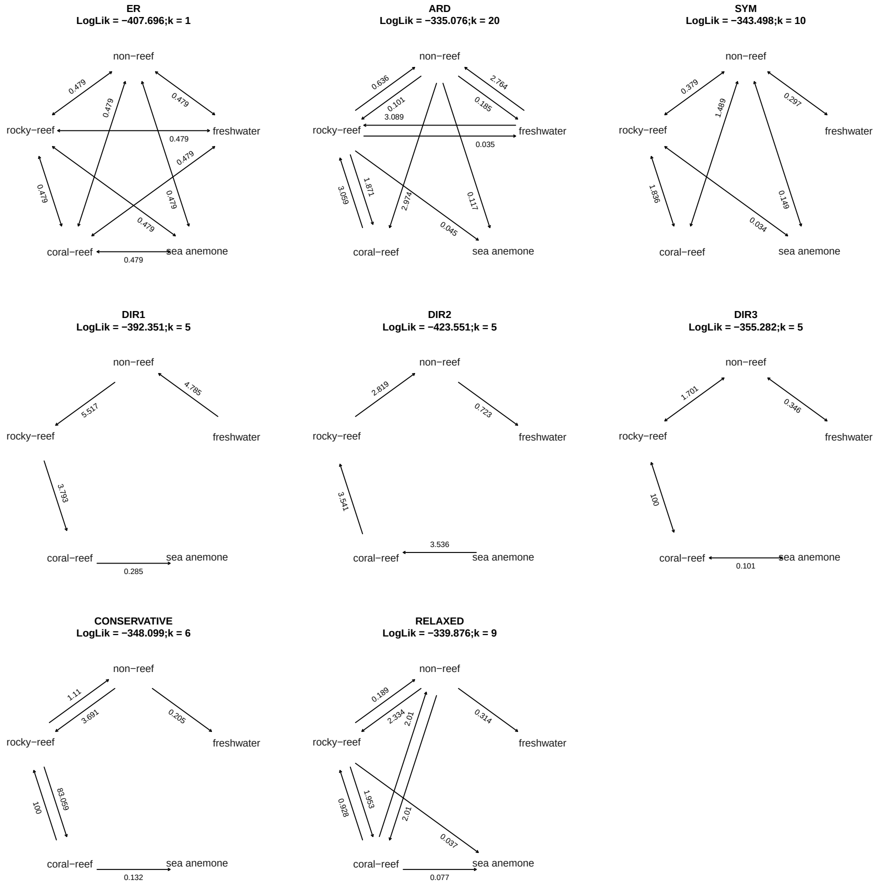

**(c) Symbiosis**

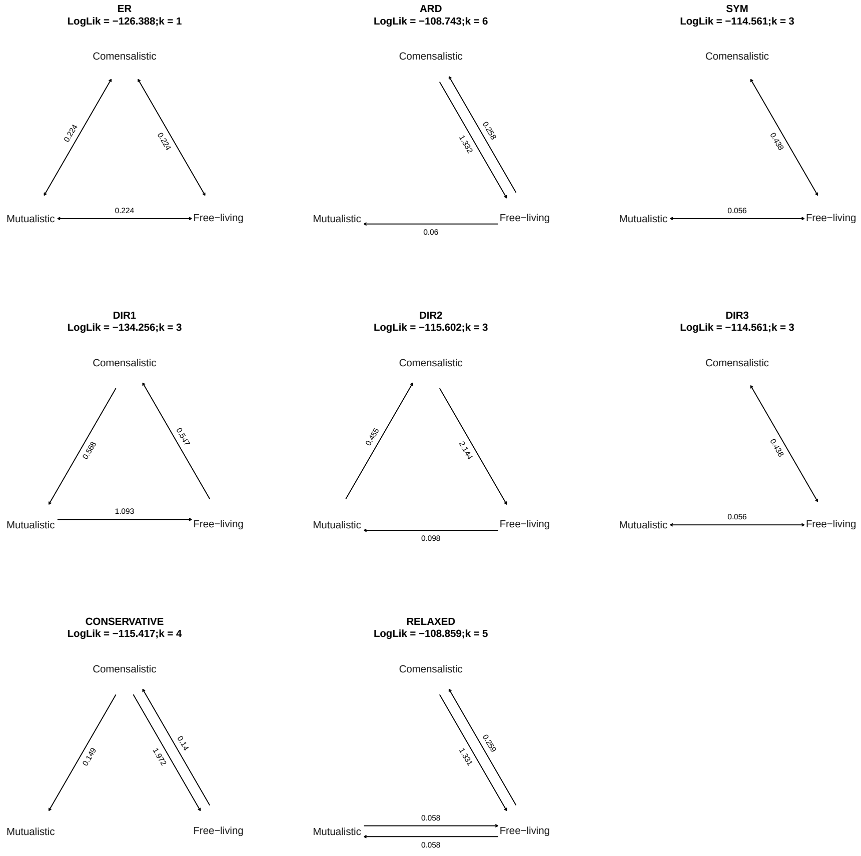


**Fig. 10**: Fitted Mk model transition-rate matrices ($Q$) for (a) diet ecotype, (b) habitat type, and (c) symbiosis. Each panel shows transition rates under alternative configurations, including equal rates (ER), all rates different (ARD), symmetric (SYM), and directionally constrained models (DIR1–DIR3). Additionally, biologically informed transition matrices ("CONSERVATIVE" and "RELAXED") were implemented for each trait to reflect empirically supported evolutionary constraints. For *DietEcotype* and *Symbiosis*, direct transitions between extreme states (e.g., benthic $\leftrightarrow$ pelagic or free-living $\leftrightarrow$ mutualistic) were restricted in the conservative model and allowed in the relaxed model. For *Habitat*, the matrix captured the hierarchical and asymmetric nature of habitat evolution in Pomacentridae: stepwise transitions among marine habitats (non-reef $\leftrightarrow$ rocky-reef $\leftrightarrow$ coral-reef) were permitted, whereas direct shifts between freshwater and fully marine or symbiotic habitats were constrained. These directional restrictions reflect the well-supported marine origin of damselfishes [9] and the extremely low probability of secondary colonisation of marine habitats from freshwater lineages [10]. Log-likelihood (*LogLik*) and the number of estimated parameters ($k$) are indicated at the top of each panel.

#### (a) Diet ecotype

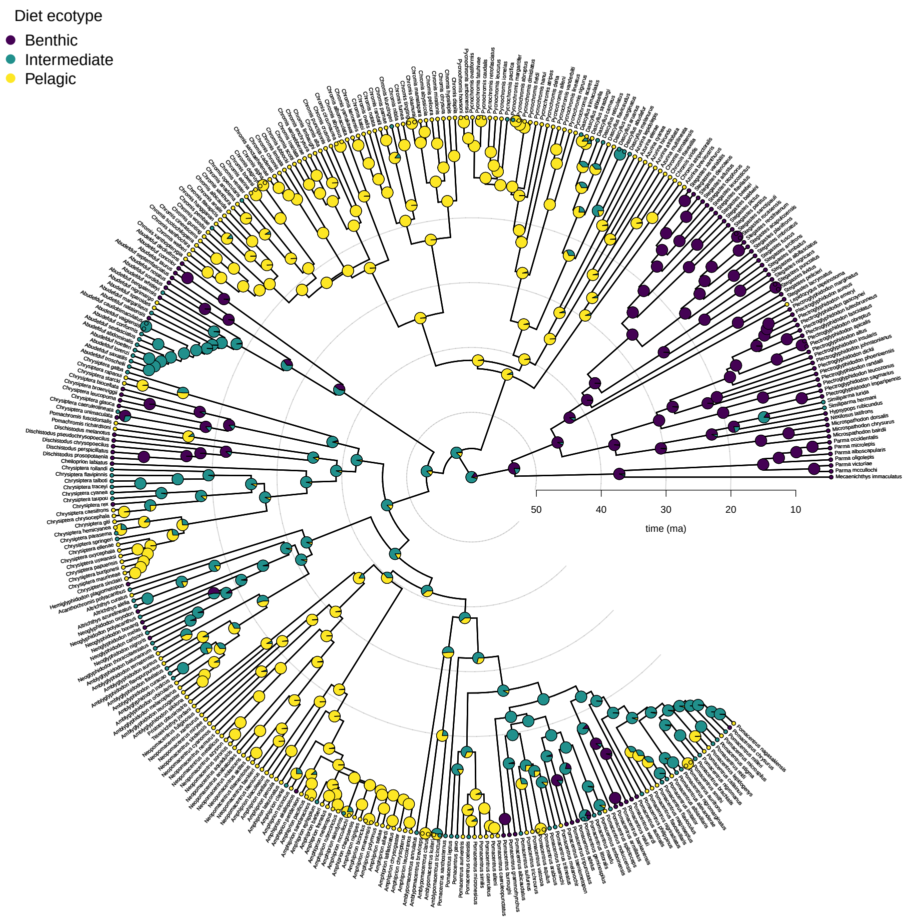


#### (b) Habitat type

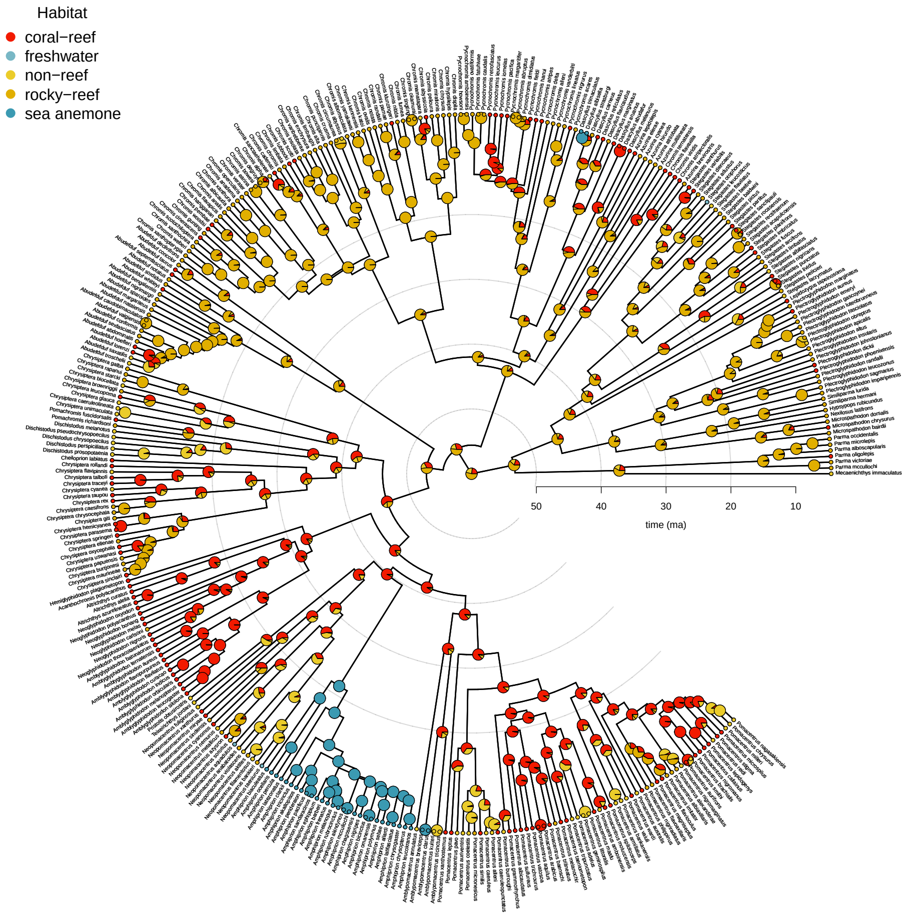

**(c) Symbiosis**

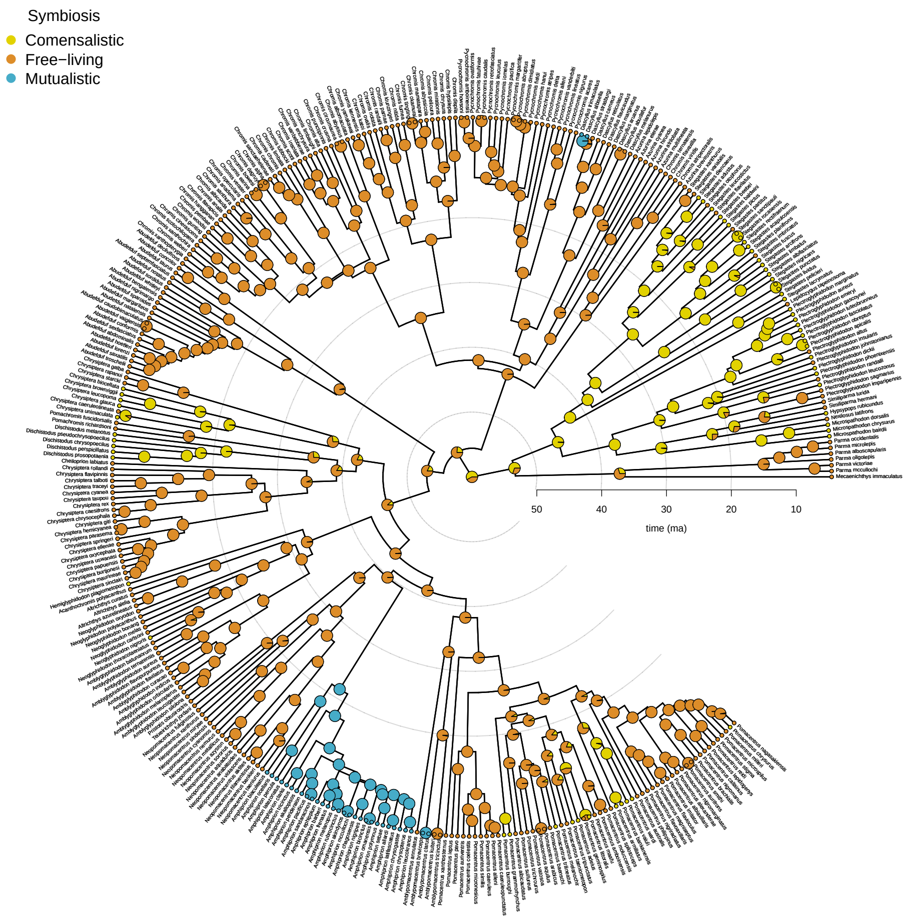


**Fig. 11**: Ancestral state reconstructions of dietary ecotype, habitat type and symbiotic relationships of the Pomacentridae family. The CONSERVATIVE model was selected for diet ecotype while the RELAXED model was selected for habitat and symbiosis, following AICc criterion [11], to fit their evolutionary trajectories. Coloured circles at the tips display the current dietary strategy (a) and habitat preference (b) and symbiosis (c). Coloured pies at the nodes represent the proportion of 1000 stochastic maps of the ancestral state reconstruction agreeing with a specific dietary strategy (a), habitat preference (b) or symbiosis (c) assignment. Colour-code legends representing each dietary strategy or habitat preference are shown at the top-left of each plot.

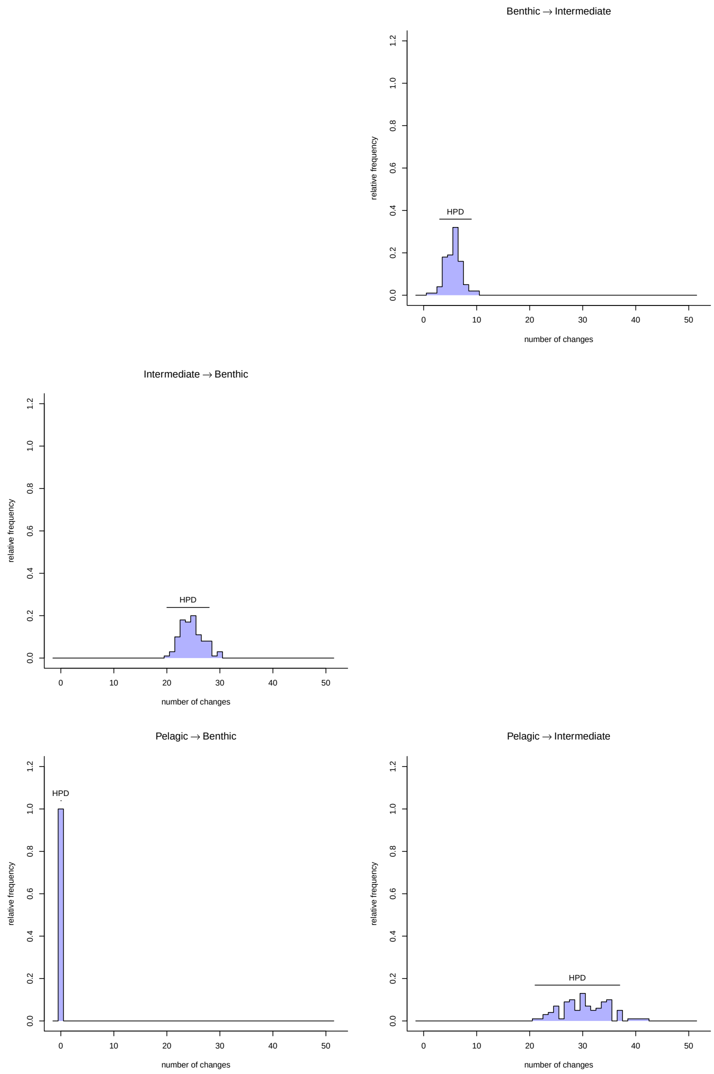


**Fig. 12**: Histograms of event counts in each of the 1000 ancestral stochastic maps (ASM) carried out on diet ecotype following the CONSERVATIVE transition model. For each type of transition, the y axis indicates the number of ASM with the same number of counts for that transition, while x axis indicates the number of counts of the particular transition.

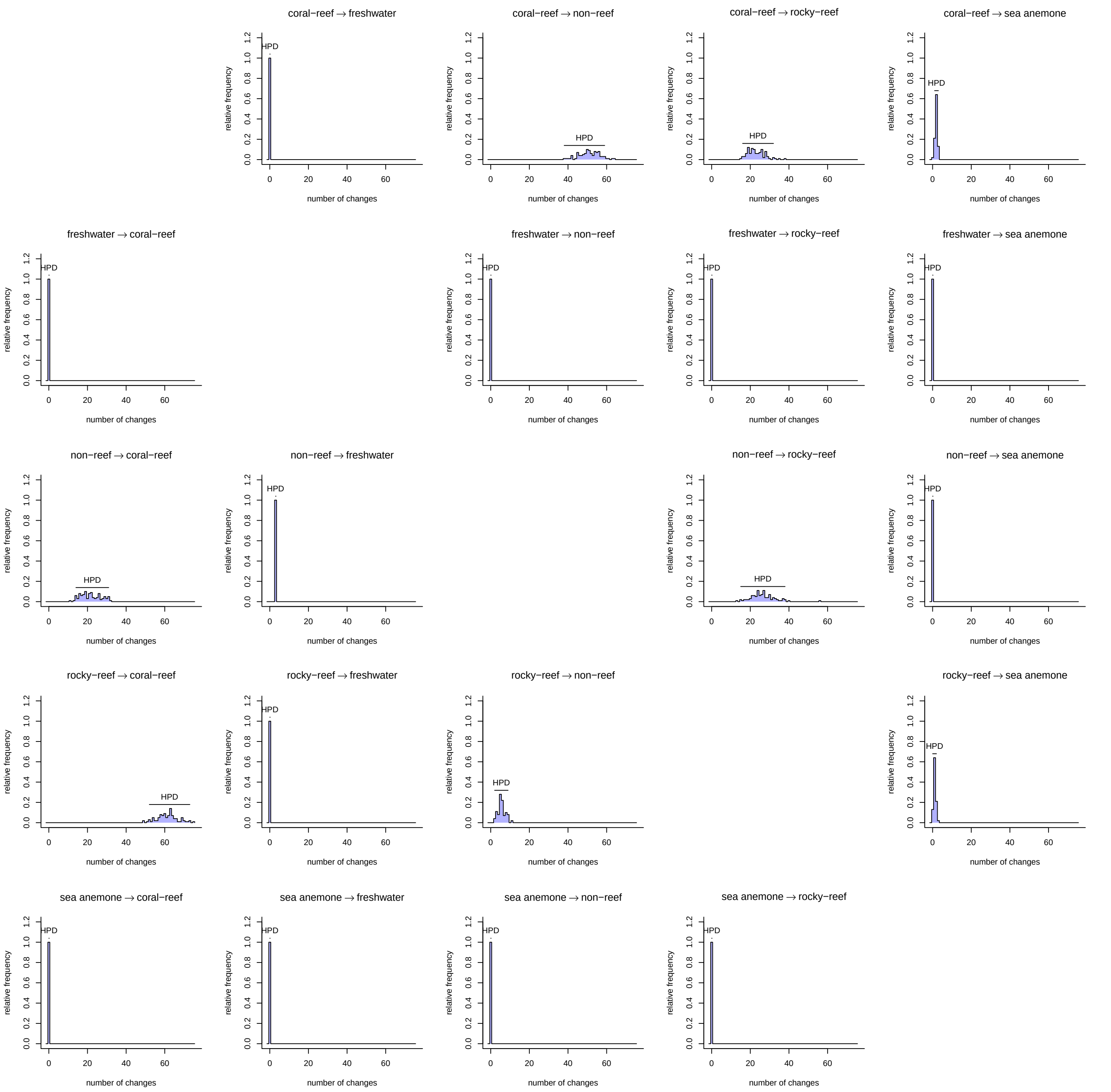


**Fig. 13**: Histograms of event counts in each of the 1000 ancestral stochastic maps (ASM) carried out on habitat type following the RELAXED transition model. For each type of transition, the y axis indicates the number of ASM with the same number of counts for that transition, while x axis indicates the number of counts of the particular transition.

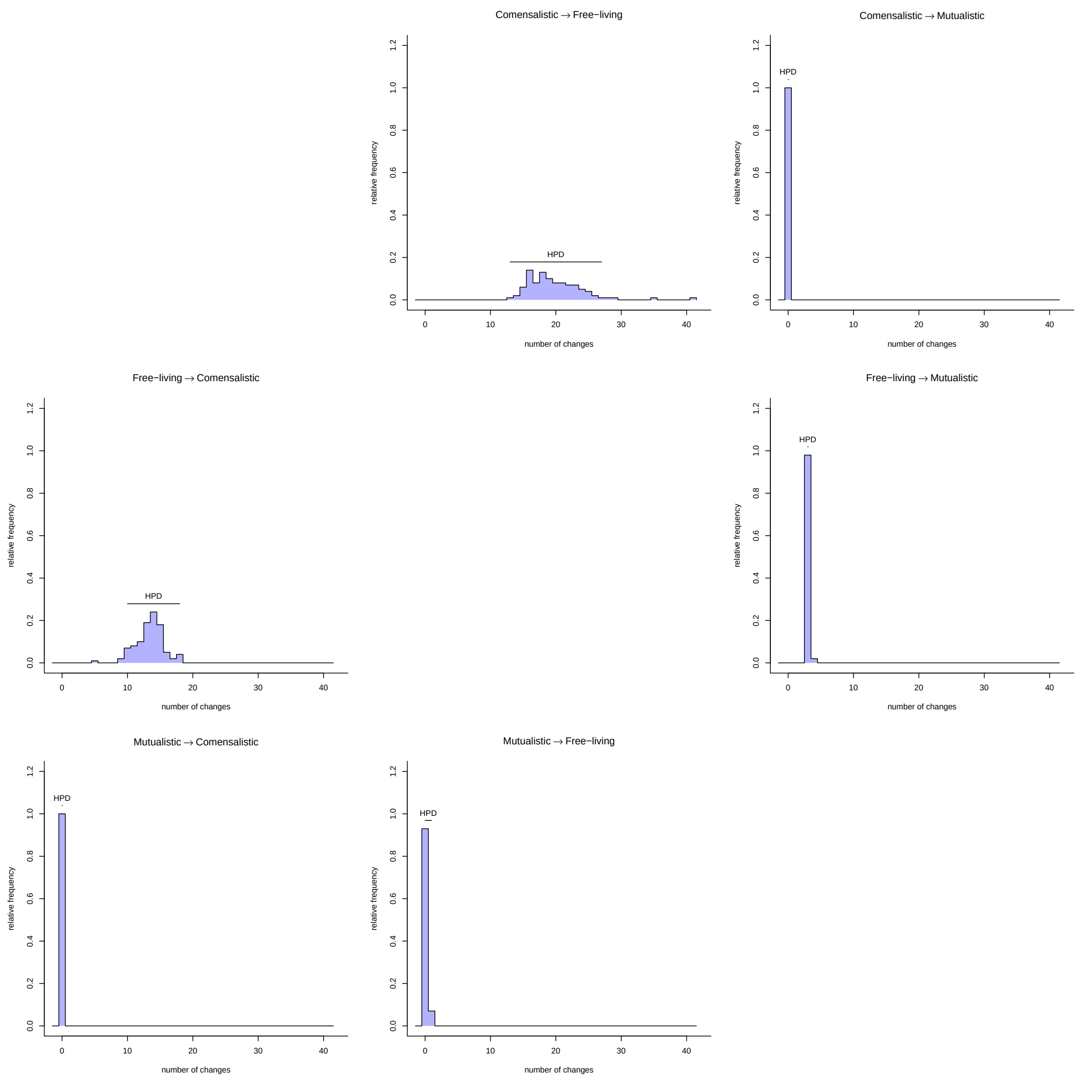


**Fig. 14**: Histograms of event counts in each of the 1000 ancestral stochastic maps (ASM) carried out on symbiotic relationships following the RELAXED transition model. For each type of transition, the y axis indicates the number of ASM with the same number of counts for that transition, while x axis indicates the number of counts of the particular transition.

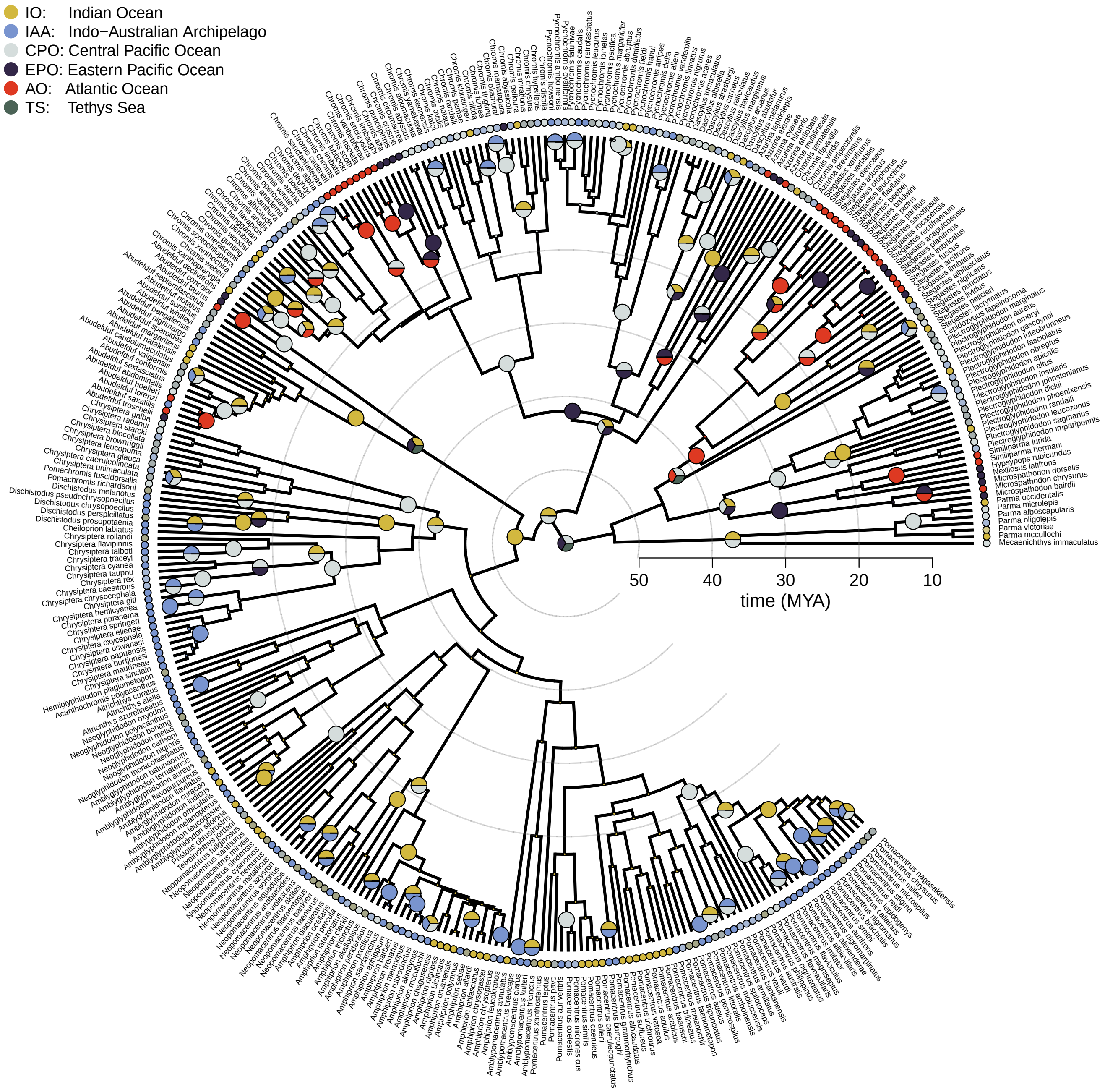


**Fig. 15**: Ancestral biogeographic reconstruction of the Pomacentridae family. The best-fitting model based on AICc was "DEC+J", a Dispersal–Extinction–Cladogenesis model incorporating founder-event speciation (+J). Coloured circles at the tips indicate present-day bioregions, while circles at the nodes represent inferred ancestral ranges; multicoloured nodes denote occupancy of multiple regions. For clarity, only nodes associated with range transitions (colonisation or expansion events) are displayed. The colour legend for each bioregion is shown in the top left. The horizontal scale represents time in millions of years before present (Mya).

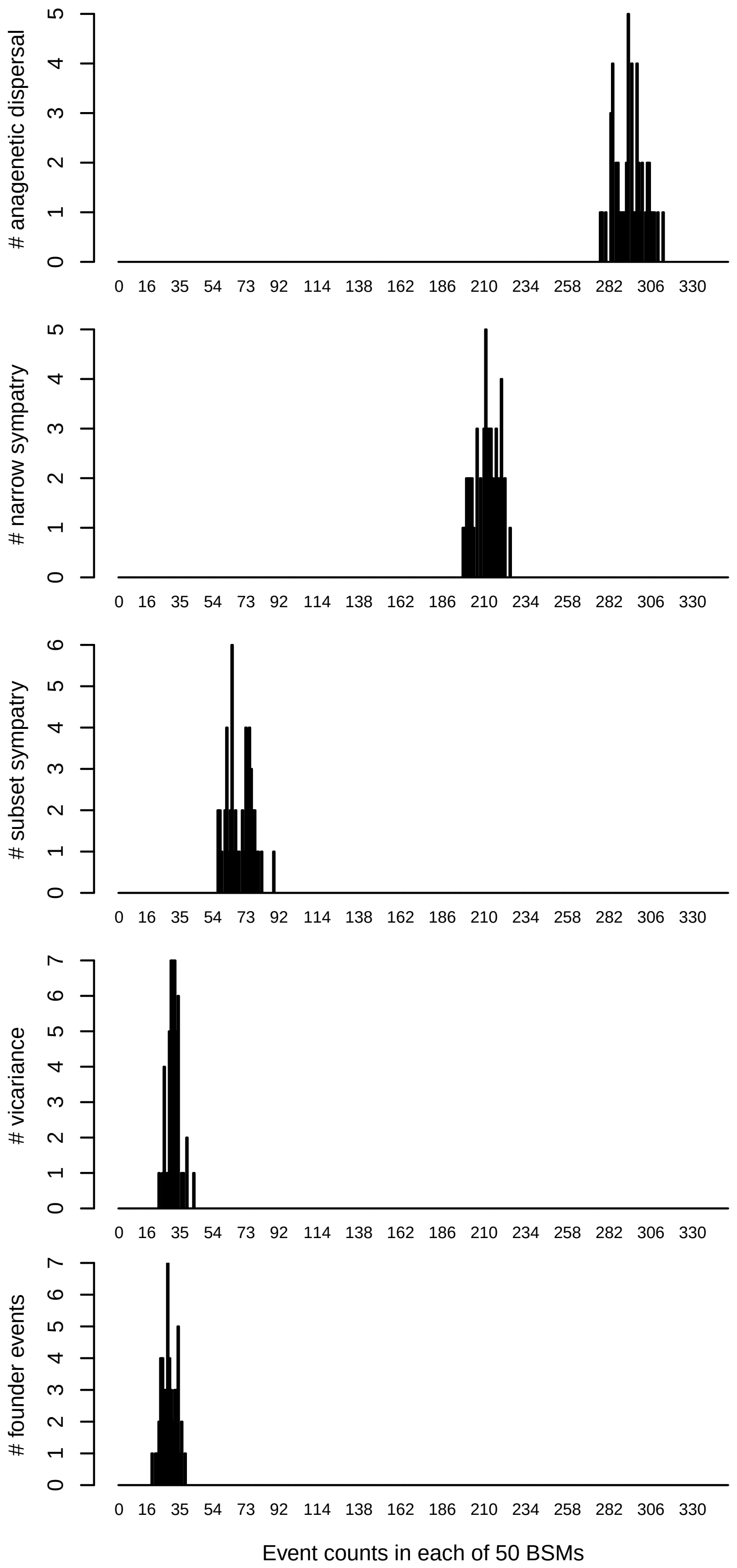


**Fig. 16**: Histogram of event counts in each of the 50 BSMs carried out. For each event, the y axis indicates the number of BSMs with the same number of counts for that event, while x axis indicates the number of counts of the particular event.

#### Supplementary Tables

**Table 1**: Summary of all ℓ1ou regime shifts for colour PCs, including shift magnitudes from ancestor's optimum, inferred regimes, bootstrap support, and qualitative trait patterns. Negative values represent shifts toward negative PC scores.

| PC | Clade(s) | Shift Value | Regime | Bootstrap | Colour Pattern |
|---|---|---|---|---|---|
| PC2 | *Amphiprion* | 194.38 | R1 | 0.97 | High contrast; white bars; segmentation. |
| | Microspathodontinae | 86.46 | R2 | 0.70 | Dorsolateral contrast; edge patterning. |
| | *Abudefduf* | –114.74 | R3 | 0.33 | Darker body; low contrast. |
| | *C. cyanea, C. taupou* | –322.31 | R4 | 0.79 | Uniform dark blue. |
| PC3 | *Dascyllus* | –98.86 | R1 | 0.58 | Darker posterior; reduced gradient. |
| PC4 | *Amphiprion* | –110.92 | R1 | 0.98 | Darker mid□body; fewer markings. |
| | *Dascyllus* | –119.45 | R1 | 0.81 | Darker overall. |
| | *Hypsypops* | –216.32 | R2 | 0.65 | Orange□red saturation. |
| | *Chrysiptera, Pomacentrus* | –156.13 to –156.98 | R3 | 0.87–0.84 | Reduced striping; dark patches. |
| PC5 | *Pycnochromis* | 159.04 | R1 | 0.53 | Brightening; highlight patches. |
| PC6 | *Pycnochromis* | 155.97 | R1 | 0.79 | Brighter ventral region. |
| | *Chrysiptera, Dascyllus, Pomacentrus* | –119.29 to –134.60 | R2 | 0.41–0.93 | Duller pigmentation. |
| | *Amphiprion* (broad) | –96.01 | R3 | 0.82 | Darker body. |
| | *Amphiprion* (Skunk complex) | 126.21 | Baseline | 0.54 | Pale dorsal stripe. |
| PC7 | *Amblypomacentrus, Amphiprion* | –88.6 to –97.45 | R1 | 0.54–0.91 | Dark head; reduced contrast. |
| | *Amblypomacentrus, Dascyllus, A. clarkii, A. latifasciatus* | –234.43 to –271.81 | R2 | 0.46–1.00 | Strong uniform darkening. |
| | *Dascyllus + Pycnochromis* | –34.73 | R3 | 0.54 | Subtle darkening. |
| | *Similiparma hermani* | –141.85 | R4 | 0.59 | Dorsal darkening. |
| | *A. latezonatus* | –303.42 | R5 | 1.00 | Extreme darkening. |
| PC8 | *Hypsypops rubicundus* | –186 | R1 | 0.61 | Orange□red saturation; single shift. |

**Table 2**: Summary of all $\ell$1ou regime shifts for morphology PCs, including shift values from ancestor's optimum, inferred regimes, bootstrap support, and associated morphological patterns. Negative values indicate shifts toward negative PC scores.

| PC | Clade(s) | Shift Value | Regime | Bootstrap | Morphological Pattern |
|---|---|---|---|---|---|
| PC1 | *Chromis* (subclades 1–2) | 21.10, 18.10 | R1 | 0.27–0.41 | Elongated, streamlined body. |
| | *Azurina*, *Neopomacentrus*, *Pristotis obtusirostris*, *Teixeirichthys jordani* | 23.99–28.05 | R2 | 0.54–0.93 | Midwater elongation. |
| | *Cheiloprion*, *Chrysiptera*, *Dischistodus*, *Pomachromis*; *Chromis* (subclade 3) | 9.20–10.77 | R3 | 0.14–0.71 | Mild elongation. |
| | *Pycnochromis* | 38.96 | R4 | 0.92 | Very slender pelagic form. |
| PC2 | *Chrysiptera* + *Cheiloprion* | –13.06 | R1 | 0.70 | Deeper body profile. |
| PC3 | *Amblyglyphidodon* | –9.04 | R1 | 0.68 | Shorter snout. |
| | *Dischistodus* | 11.10 | R2 | 0.40 | Robust anterior. |
| | *Azurina*, *Chromis*, *Dascyllus*, *Pycnochromis* | –4.28 | R3 | 0.52 | Reduced head depth. |
| | *Dascyllus*, *Pycnochromis* | –8.83 | R4 | 0.96 | Thickened head. |
| PC4 | *Amblyglyphidodon*, *Dascyllus* | –12.90 to –13.17 | R1 | 0.90–0.97 | Deeper mid□body. |
| | *Abudefduf* | –9.38 | R2 | 0.51 | Slight increase in depth. |
| | *Pomacentrus* | 4.29 | R3 | 0.36 | Narrower mid□body. |
| PC5 | *Amphiprion*, *Neopomacentrus*, *Pristotis*, *Teixeirichthys* | –4.54 | R1 | 0.84 | Deeper anterior body. |
| | *Chrysiptera flavipinnis*, *C. rollandi*, *C. talboti*, *C. traceyi* | 11.30 | R2 | 0.50 | Slimmer head. |
| PC6 | *Stegastes* | –6.05 | R1 | 0.46 | Short snout. |
| | *Pycnochromis*, Pomacentrinae | –7.73 to –9.14 | R2 | 0.72–1.00 | Slimmer body; short snout. |
| | *Abudefduf* | 5.67 | R3 | 0.80 | Deeper head. |
| | Multiple lineages | –10.80 to –21.28 | R4 | 0.64–0.96 | Strong body compression. |
| | *Chrysiptera rollandi* | 19.51 | R5 | 0.74 | Deeper body/head. |
| | *Pomacentrus australis* | –17.94 | R6 | 0.66 | Body compression. |
| | *Acanthochromis*, *Altrichthys*, *Neoglyphidodon*, *Dischistodus* | 8.97–9.34 | R7 | 0.76–0.94 | Reversal toward ancestral. |
| PC7 | *Lepidozygus tapeinosoma* | –17.29 | R1 | 0.65 | Slender body. |
| | *Azurina* spp. | 8.98 | R2 | 0.26 | Robust head. |
| | Chrominae, Microspathodontinae | 3.91–5.18 | R3 | 0.55–0.58 | Slight increase in depth. |
| PC8 | *Amphiprion bicinctus*, *A. omanensis* | 11.99 | R1 | 0.78 | Robust snout. |
| | *Chrysiptera rollandi* | 16.25 | R2 | 0.94 | Head enlargement. |
| | *Chromis cadenati* | –13.95 | R3 | 0.70 | Slender head. |

**Table 3**: Comparison of alternative phylogenetic correlation structures (BM, Pagel's $\lambda$, OU) in univariate PGLS models of colour PCs, using the best ecological predictor for each PC. The reported parameter is $\lambda$ for Pagel's model and $\alpha$ for OU (BM has no parameter). Within each PC, the best-fitting structure (lowest AICc) is shown in bold.

| PC | Predictor | Correlation | Param | df | AICc | ΔAICc | Weight |
|---|---|---|---|---|---|---|---|
| | **NULL** | **Pagel ($\lambda$)** | **0.185** | 3 | **973.737** | **0.000** | **0.976** |
| PC1 | NULL | OU ($\alpha$) | 2.523 | 3 | 981.120 | 7.383 | 0.024 |
| | NULL | BM | — | 2 | 1305.905 | 332.168 | 0.000 |
| | **NULL** | **Pagel ($\lambda$)** | **0.266** | 3 | **951.838** | **0.000** | **1.000** |
| PC2 | NULL | OU ($\alpha$) | 1.579 | 3 | 979.488 | 27.650 | 0.000 |
| | NULL | BM | — | 2 | 1286.788 | 334.951 | 0.000 |
| | **NULL** | **Pagel ($\lambda$)** | **0.079** | 3 | **979.552** | **0.000** | **0.615** |
| PC3 | NULL | OU ($\alpha$) | 4.795 | 3 | 980.486 | 0.934 | 0.385 |
| | NULL | BM | — | 2 | 1427.964 | 448.412 | 0.000 |
| | **Habitat+Diet** | **Pagel ($\lambda$)** | **-0.080** | 9 | **960.421** | **0.000** | **1.000** |
| PC4 | Habitat+Diet | OU ($\alpha$) | 15.992 | 9 | 990.832 | 30.411 | 0.000 |
| | Habitat+Diet | BM | — | 8 | 1391.199 | 430.778 | 0.000 |
| | **NULL** | **OU ($\alpha$)** | **1.044** | 3 | **976.725** | **0.000** | **0.504** |
| PC5 | NULL | Pagel ($\lambda$) | 0.159 | 3 | 976.754 | 0.029 | 0.496 |
| | NULL | BM | — | 2 | 1249.585 | 272.861 | 0.000 |
| | **NULL** | **Pagel ($\lambda$)** | **0.236** | 3 | **952.397** | **0.000** | **1.000** |
| PC6 | NULL | OU ($\alpha$) | 2.042 | 3 | 980.192 | 27.795 | 0.000 |
| | NULL | BM | — | 2 | 1281.880 | 329.483 | 0.000 |
| | **NULL** | **Pagel ($\lambda$)** | **0.368** | 3 | **952.955** | **0.000** | **1.000** |
| PC7 | NULL | OU ($\alpha$) | 2.482 | 3 | 981.421 | 28.466 | 0.000 |
| | NULL | BM | — | 2 | 1292.468 | 339.513 | 0.000 |
| | **NULL** | **Pagel ($\lambda$)** | **0.179** | 3 | **980.473** | **0.000** | **0.540** |
| PC8 | NULL | OU ($\alpha$) | 3.317 | 3 | 980.792 | 0.320 | 0.460 |
| | NULL | BM | — | 2 | 1348.282 | 367.809 | 0.000 |

**Table 4**: Comparison of alternative phylogenetic correlation structures (BM, Pagel's $\lambda$, OU) in univariate PGLS models of morphology PCs, using the best ecological predictor for each PC. The reported parameter is $\lambda$ for Pagel's model and $\alpha$ for OU (BM has no parameter). Within each PC, the best-fitting structure (lowest AICc) is shown in bold.

| PC | Predictor | Correlation | Param | df | AICc | ΔAICc | Weight |
|---|---|---|---|---|---|---|---|
| PC1 | **NULL** | **Pagel ($\lambda$)** | **0.065** | 3 | **974.344** | **0.000** | **0.983** |
| | NULL | OU ($\alpha$) | 61.774 | 3 | 982.462 | 8.118 | 0.017 |
| | NULL | BM | — | 2 | 1514.176 | 539.832 | 0.000 |
| PC2 | **NULL** | **Pagel ($\lambda$)** | **0.316** | 3 | **949.682** | **0.000** | **1.000** |
| | NULL | OU ($\alpha$) | 1.235 | 3 | 977.650 | 27.968 | 0.000 |
| | NULL | BM | — | 2 | 1292.547 | 342.865 | 0.000 |
| PC3 | **NULL** | **Pagel ($\lambda$)** | **0.310** | 3 | **948.829** | **0.000** | **1.000** |
| | NULL | OU ($\alpha$) | 6.044 | 3 | 982.316 | 33.487 | 0.000 |
| | NULL | BM | — | 2 | 1324.592 | 375.763 | 0.000 |
| PC4 | **NULL** | **Pagel ($\lambda$)** | **0.392** | 3 | **951.027** | **0.000** | **1.000** |
| | NULL | OU ($\alpha$) | 0.946 | 3 | 974.357 | 23.330 | 0.000 |
| | NULL | BM | — | 2 | 1267.227 | 316.199 | 0.000 |
| PC5 | **DietEcotype** | **Pagel ($\lambda$)** | **0.115** | 5 | **964.191** | **0.000** | **0.994** |
| | DietEcotype | OU ($\alpha$) | 3.237 | 5 | 974.272 | 10.080 | 0.006 |
| | DietEcotype | BM | — | 4 | 1354.133 | 389.942 | 0.000 |
| PC6 | **NULL** | **Pagel ($\lambda$)** | **0.628** | 3 | **818.603** | **0.000** | **1.000** |
| | NULL | OU ($\alpha$) | 0.684 | 3 | 974.721 | 156.118 | 0.000 |
| | NULL | BM | — | 2 | 1198.723 | 380.120 | 0.000 |
| PC7 | **NULL** | **Pagel ($\lambda$)** | **0.266** | 3 | **950.733** | **0.000** | **1.000** |
| | NULL | OU ($\alpha$) | 1.256 | 3 | 977.416 | 26.684 | 0.000 |
| | NULL | BM | — | 2 | 1271.807 | 321.074 | 0.000 |
| PC8 | **NULL** | **Pagel ($\lambda$)** | **0.139** | 3 | **970.545** | **0.000** | **0.997** |
| | NULL | OU ($\alpha$) | 4.626 | 3 | 981.921 | 11.375 | 0.003 |
| | NULL | BM | — | 2 | 1340.876 | 370.330 | 0.000 |

**Table 5**: Comparison of ecological predictor models in univariate PGLS analyses of colour PCs, with the best-fitting correlation structure selected for each predictor. The reported parameter is $\lambda$ for Pagel's model and $\alpha$ for OU (BM has no parameter). Within each PC, the best predictor model (lowest AICc) is shown in bold.

| PC | Predictor | Correlation | Param | df | AICc | ΔAICc | Weight |
|---|---|---|---|---|---|---|---|
| PC1 | **NULL** | **Pagel ($\lambda$)** | **0.185** | 3 | **973.737** | **0.000** | **0.886** |
| | Symbiosis | Pagel ($\lambda$) | 0.203 | 5 | 978.617 | 4.880 | 0.077 |
| | DietEcotype | Pagel ($\lambda$) | 0.192 | 5 | 981.274 | 7.538 | 0.020 |
| | Habitat | Pagel ($\lambda$) | 0.186 | 7 | 982.067 | 8.330 | 0.014 |
| | Symbiosis+Diet | Pagel ($\lambda$) | 0.206 | 7 | 985.721 | 11.984 | 0.002 |
| | Habitat+Diet | Pagel ($\lambda$) | 0.192 | 9 | 989.722 | 15.985 | 0.000 |
| PC2 | **NULL** | **Pagel ($\lambda$)** | **0.266** | 3 | **951.838** | **0.000** | **0.889** |
| | DietEcotype | Pagel ($\lambda$) | 0.253 | 5 | 957.656 | 5.818 | 0.048 |
| | Symbiosis | Pagel ($\lambda$) | 0.279 | 5 | 957.795 | 5.957 | 0.045 |
| | Habitat | Pagel ($\lambda$) | 0.289 | 7 | 960.246 | 8.408 | 0.013 |
| | Symbiosis+Diet | Pagel ($\lambda$) | 0.264 | 7 | 963.332 | 11.494 | 0.003 |
| | Habitat+Diet | Pagel ($\lambda$) | 0.274 | 9 | 965.728 | 13.890 | 0.001 |
| PC3 | **NULL** | **Pagel ($\lambda$)** | **0.079** | 3 | **979.552** | **0.000** | **0.740** |
| | Symbiosis | Pagel ($\lambda$) | 0.081 | 5 | 983.014 | 3.462 | 0.131 |
| | DietEcotype | Pagel ($\lambda$) | 0.101 | 5 | 984.123 | 4.572 | 0.075 |
| | Symbiosis+Diet | Pagel ($\lambda$) | 0.064 | 7 | 985.709 | 6.157 | 0.034 |
| | Habitat | OU ($\alpha$) | 4.521 | 7 | 986.974 | 7.422 | 0.018 |
| | Habitat+Diet | OU ($\alpha$) | 4.607 | 9 | 991.155 | 11.604 | 0.002 |
| PC4 | **Habitat+Diet** | **Pagel ($\lambda$)** | **-0.080** | 9 | **960.421** | **0.000** | **0.907** |
| | Symbiosis+Diet | Pagel ($\lambda$) | -0.047 | 7 | 965.339 | 4.918 | 0.078 |
| | NULL | Pagel ($\lambda$) | 0.154 | 3 | 969.621 | 9.200 | 0.009 |
| | Symbiosis | Pagel ($\lambda$) | 0.115 | 5 | 970.544 | 10.123 | 0.006 |
| | DietEcotype | Pagel ($\lambda$) | 0.153 | 5 | 976.641 | 16.220 | 0.000 |
| | Habitat | Pagel ($\lambda$) | 0.130 | 7 | 977.379 | 16.958 | 0.000 |
| PC5 | **NULL** | **OU ($\alpha$)** | **1.044** | 3 | **976.725** | **0.000** | **0.752** |
| | Symbiosis | Pagel ($\lambda$) | 0.157 | 5 | 980.388 | 3.663 | 0.120 |
| | DietEcotype | OU ($\alpha$) | 1.041 | 5 | 981.189 | 4.465 | 0.081 |
| | Habitat | OU ($\alpha$) | 1.082 | 7 | 983.388 | 6.663 | 0.027 |
| | Symbiosis+Diet | OU ($\alpha$) | 1.107 | 7 | 984.438 | 7.714 | 0.016 |
| | Habitat+Diet | OU ($\alpha$) | 1.142 | 9 | 987.041 | 10.317 | 0.004 |
| PC6 | **NULL** | **Pagel ($\lambda$)** | **0.236** | 3 | **952.397** | **0.000** | **0.280** |
| | Habitat | Pagel ($\lambda$) | 0.260 | 7 | 952.637 | 0.240 | 0.249 |
| | DietEcotype | Pagel ($\lambda$) | 0.266 | 5 | 953.263 | 0.865 | 0.182 |
| | Habitat+Diet | Pagel ($\lambda$) | 0.277 | 9 | 953.463 | 1.066 | 0.165 |
| | Symbiosis | Pagel ($\lambda$) | 0.233 | 5 | 954.580 | 2.182 | 0.094 |
| | Symbiosis+Diet | Pagel ($\lambda$) | 0.249 | 7 | 956.840 | 4.443 | 0.030 |
| PC7 | **NULL** | **Pagel ($\lambda$)** | **0.368** | 3 | **952.955** | **0.000** | **0.845** |
| | Symbiosis | Pagel ($\lambda$) | 0.370 | 5 | 957.706 | 4.751 | 0.079 |
| | DietEcotype | Pagel ($\lambda$) | 0.394 | 5 | 958.765 | 5.810 | 0.046 |
| | Symbiosis+Diet | Pagel ($\lambda$) | 0.382 | 7 | 960.113 | 7.158 | 0.024 |
| | Habitat | Pagel ($\lambda$) | 0.382 | 7 | 962.848 | 9.893 | 0.006 |
| | Habitat+Diet | Pagel ($\lambda$) | 0.414 | 9 | 968.113 | 15.158 | 0.000 |
| PC8 | **NULL** | **Pagel ($\lambda$)** | **0.179** | 3 | **980.473** | **0.000** | **0.825** |
| | Symbiosis | Pagel ($\lambda$) | 0.203 | 5 | 984.225 | 3.752 | 0.126 |
| | DietEcotype | Pagel ($\lambda$) | 0.159 | 5 | 986.902 | 6.429 | 0.033 |
| | Habitat | Pagel ($\lambda$) | 0.262 | 7 | 989.231 | 8.758 | 0.010 |
| | Symbiosis+Diet | Pagel ($\lambda$) | 0.201 | 7 | 991.248 | 10.775 | 0.004 |
| | Habitat+Diet | Pagel ($\lambda$) | -0.138 | 9 | 993.396 | 12.923 | 0.001 |

**Table 6**: Comparison of ecological predictor models in univariate PGLS analyses of morphology PCs, with the best-fitting correlation structure selected for each predictor. The reported parameter is $\lambda$ for Pagel's model and $\alpha$ for OU (BM has no parameter). Within each PC, the best predictor model (lowest AICc) is shown in bold.

| PC | Predictor | Correlation | Param | df | AICc | ΔAICc | Weight |
|---|---|---|---|---|---|---|---|
| PC1 | **NULL** | **Pagel ($\lambda$)** | **0.065** | 3 | **974.344** | **0.000** | **0.797** |
| | DietEcotype | Pagel ($\lambda$) | 0.063 | 5 | 978.172 | 3.827 | 0.118 |
| | Symbiosis | Pagel ($\lambda$) | 0.074 | 5 | 980.160 | 5.816 | 0.044 |
| | Symbiosis+Diet | Pagel ($\lambda$) | -0.084 | 7 | 980.887 | 6.543 | 0.030 |
| | Habitat+Diet | Pagel ($\lambda$) | -0.083 | 9 | 983.416 | 9.072 | 0.009 |
| | Habitat | Pagel ($\lambda$) | 0.062 | 7 | 985.462 | 11.117 | 0.003 |
| PC2 | **NULL** | **Pagel ($\lambda$)** | **0.316** | 3 | **949.682** | **0.000** | **0.655** |
| | Habitat | Pagel ($\lambda$) | 0.291 | 7 | 951.456 | 1.774 | 0.270 |
| | Symbiosis | Pagel ($\lambda$) | 0.315 | 5 | 954.913 | 5.230 | 0.048 |
| | DietEcotype | Pagel ($\lambda$) | 0.324 | 5 | 957.045 | 7.363 | 0.017 |
| | Habitat+Diet | Pagel ($\lambda$) | 0.298 | 9 | 958.260 | 8.578 | 0.009 |
| | Symbiosis+Diet | Pagel ($\lambda$) | 0.322 | 7 | 961.805 | 12.122 | 0.002 |
| PC3 | **NULL** | **Pagel ($\lambda$)** | **0.310** | 3 | **948.829** | **0.000** | **0.818** |
| | DietEcotype | Pagel ($\lambda$) | 0.281 | 5 | 953.147 | 4.317 | 0.095 |
| | Symbiosis | Pagel ($\lambda$) | 0.319 | 5 | 954.274 | 5.444 | 0.054 |
| | Habitat | Pagel ($\lambda$) | 0.315 | 7 | 956.303 | 7.473 | 0.020 |
| | Symbiosis+Diet | Pagel ($\lambda$) | 0.275 | 7 | 957.233 | 8.403 | 0.012 |
| | Habitat+Diet | Pagel ($\lambda$) | 0.291 | 9 | 961.527 | 12.698 | 0.001 |
| PC4 | **NULL** | **Pagel ($\lambda$)** | **0.392** | 3 | **951.027** | **0.000** | **0.898** |
| | Symbiosis | Pagel ($\lambda$) | 0.400 | 5 | 956.748 | 5.721 | 0.051 |
| | DietEcotype | Pagel ($\lambda$) | 0.404 | 5 | 957.256 | 6.229 | 0.040 |
| | Habitat | Pagel ($\lambda$) | 0.402 | 7 | 960.537 | 9.509 | 0.008 |
| | Symbiosis+Diet | Pagel ($\lambda$) | 0.410 | 7 | 962.377 | 11.350 | 0.003 |
| | Habitat+Diet | Pagel ($\lambda$) | 0.413 | 9 | 966.438 | 15.410 | 0.000 |
| PC5 | **DietEcotype** | **Pagel ($\lambda$)** | **0.115** | 5 | **964.191** | **0.000** | **0.555** |
| | NULL | Pagel ($\lambda$) | 0.139 | 3 | 966.006 | 1.815 | 0.224 |
| | Habitat | Pagel ($\lambda$) | 0.115 | 7 | 967.465 | 3.274 | 0.108 |
| | Habitat+Diet | Pagel ($\lambda$) | 0.097 | 9 | 968.320 | 4.129 | 0.070 |
| | Symbiosis+Diet | Pagel ($\lambda$) | 0.121 | 7 | 969.883 | 5.691 | 0.032 |
| | Symbiosis | Pagel ($\lambda$) | 0.147 | 5 | 972.180 | 7.989 | 0.010 |
| PC6 | **NULL** | **Pagel ($\lambda$)** | **0.628** | 3 | **818.603** | **0.000** | **0.808** |
| | Symbiosis | Pagel ($\lambda$) | 0.618 | 5 | 822.194 | 3.591 | 0.134 |
| | Habitat | Pagel ($\lambda$) | 0.633 | 7 | 824.820 | 6.217 | 0.036 |
| | DietEcotype | Pagel ($\lambda$) | 0.625 | 5 | 826.192 | 7.589 | 0.018 |
| | Symbiosis+Diet | Pagel ($\lambda$) | 0.616 | 7 | 829.650 | 11.048 | 0.003 |
| | Habitat+Diet | Pagel ($\lambda$) | 0.630 | 9 | 832.739 | 14.136 | 0.001 |
| PC7 | **NULL** | **Pagel ($\lambda$)** | **0.266** | 3 | **950.733** | **0.000** | **0.842** |
| | Symbiosis | Pagel ($\lambda$) | 0.267 | 5 | 954.548 | 3.815 | 0.125 |
| | DietEcotype | Pagel ($\lambda$) | 0.257 | 5 | 958.043 | 7.310 | 0.022 |
| | Habitat | Pagel ($\lambda$) | 0.263 | 7 | 960.267 | 9.535 | 0.007 |
| | Symbiosis+Diet | Pagel ($\lambda$) | 0.264 | 7 | 961.198 | 10.465 | 0.004 |
| | Habitat+Diet | Pagel ($\lambda$) | 0.257 | 9 | 967.601 | 16.869 | 0.000 |
| PC8 | **NULL** | **Pagel ($\lambda$)** | **0.139** | 3 | **970.545** | **0.000** | **0.605** |
| | Symbiosis | Pagel ($\lambda$) | 0.133 | 5 | 972.464 | 1.918 | 0.232 |
| | Symbiosis+Diet | Pagel ($\lambda$) | 0.117 | 7 | 974.347 | 3.802 | 0.090 |
| | DietEcotype | Pagel ($\lambda$) | 0.137 | 5 | 975.764 | 5.218 | 0.045 |
| | Habitat | Pagel ($\lambda$) | 0.123 | 7 | 976.927 | 6.381 | 0.025 |
| | Habitat+Diet | Pagel ($\lambda$) | 0.101 | 9 | 980.681 | 10.135 | 0.004 |

**Table 7**: Univariate phylogenetic generalized least squares (PGLS) results for colour principal components (PCs). For each PC, the best-fitting ecological predictor and correlation structure (Pagel's $\lambda$ or OU with $\alpha$) were selected based on AICc. Reported are regression coefficients (Estimate), standard errors (SE), 95% confidence intervals (CIs), $t$-statistics, raw $p$-values, and false discovery rate (FDR) adjusted $p$-values. Intercepts are shown with raw $p$-values only, as they do not represent ecological contrasts.

| PC | Model | Term | Estimate | SE | 95% CI | t | p | p(FDR) |
|---|---|---|---|---|---|---|---|---|
| PC1 | NULL \| Pagel ($\lambda$=0.185) | (Intercept) | -0.064 | 0.169 | [-0.396, 0.269] | -0.38 | 0.706 | — |
| PC2 | NULL \| Pagel ($\lambda$=0.266) | (Intercept) | -0.115 | 0.196 | [-0.500, 0.271] | -0.59 | 0.558 | — |
| PC3 | NULL \| Pagel ($\lambda$=0.079) | (Intercept) | -0.059 | 0.118 | [-0.291, 0.174] | -0.49 | 0.621 | — |
| PC4 | Habitat+Diet \| Pagel ($\lambda$=-0.080) | (Intercept) | -0.178 | 0.125 | [-0.425, 0.068] | -1.42 | 0.155 | — |
| | | coral-reef | 0.170 | 0.155 | [-0.134, 0.474] | 1.10 | 0.272 | 0.544 |
| | | rocky-reef | -0.072 | 0.148 | [-0.363, 0.219] | -0.49 | 0.626 | 0.751 |
| | | sea anemone | 0.550 | 0.179 | [0.197, 0.903] | 3.07 | 0.002* | 0.007* |
| | | freshwater | -0.315 | 0.554 | [-1.405, 0.774] | -0.57 | 0.569 | 0.751 |
| | | Benthic | 0.475 | 0.070 | [0.337, 0.613] | 6.77 | <0.001* | <0.001* |
| | | Pelagic | -0.002 | 0.050 | [-0.099, 0.096] | -0.03 | 0.974 | 0.974 |
| PC5 | NULL \| OU ($\alpha$=1.044) | (Intercept) | -0.004 | 0.056 | [-0.114, 0.105] | -0.07 | 0.942 | — |
| PC6 | NULL \| Pagel ($\lambda$=0.236) | (Intercept) | 0.042 | 0.184 | [-0.320, 0.405] | 0.23 | 0.819 | — |
| PC7 | NULL \| Pagel ($\lambda$=0.368) | (Intercept) | 0.033 | 0.236 | [-0.430, 0.497] | 0.14 | 0.887 | — |
| PC8 | NULL \| Pagel ($\lambda$=0.179) | (Intercept) | 0.005 | 0.168 | [-0.325, 0.335] | 0.03 | 0.977 | — |

**Table 8**: Univariate phylogenetic generalized least squares (PGLS) results for morphological principal components (PCs). For each PC, the best-fitting ecological predictor and correlation structure (Pagel's $\lambda$ or OU with $\alpha$) were selected based on AICc. Reported are regression coefficients (Estimate), standard errors (SE), 95% confidence intervals (CIs), $t$-statistics, raw $p$-values, and false discovery rate (FDR) adjusted $p$-values. Intercepts are shown with raw $p$-values only, as they do not represent ecological contrasts.

| PC | Model | Term | Estimate | SE | 95% CI | t | p | p(FDR) |
|---|---|---|---|---|---|---|---|---|
| PC1 | NULL \| Pagel ($\lambda$=0.065) | (Intercept) | -0.103 | 0.109 | [-0.318, 0.112] | -0.94 | 0.346 | — |
| PC2 | NULL \| Pagel ($\lambda$=0.316) | (Intercept) | 0.038 | 0.215 | [-0.384, 0.460] | 0.18 | 0.860 | — |
| PC3 | NULL \| Pagel ($\lambda$=0.310) | (Intercept) | 0.042 | 0.212 | [-0.375, 0.459] | 0.20 | 0.842 | — |
| PC4 | NULL \| Pagel ($\lambda$=0.392) | (Intercept) | -0.010 | 0.244 | [-0.490, 0.471] | -0.04 | 0.969 | — |
| PC5 | DietEcotype \| Pagel ($\lambda$=0.115) | (Intercept) | -0.100 | 0.178 | [-0.449, 0.249] | -0.56 | 0.574 | — |
| | | Benthic | 0.016 | 0.178 | [-0.334, 0.367] | 0.09 | 0.927 | 0.927 |
| | | Pelagic | 0.415 | 0.147 | [0.125, 0.704] | 2.82 | 0.005* | 0.010* |
| PC6 | NULL \| Pagel ($\lambda$=0.628) | (Intercept) | 0.052 | 0.285 | [-0.508, 0.612] | 0.18 | 0.855 | — |
| PC7 | NULL \| Pagel ($\lambda$=0.266) | (Intercept) | -0.032 | 0.195 | [-0.416, 0.353] | -0.16 | 0.872 | — |
| PC8 | NULL \| Pagel ($\lambda$=0.139) | (Intercept) | -0.088 | 0.148 | [-0.378, 0.202] | -0.60 | 0.550 | — |

**Table 9**: Model comparison of multivariate phylogenetic GLS (mvGLS) analyses testing ecological predictors of combined colour and morphology principal components. Global models (BM, Pagel's $\lambda$, OU, EB) were fitted on the phylogeny, while regime-dependent models (BMM: multiple-rate BM; OUM: multiple-optima OU) were fitted on a consensus stochastic character map (SCM) for each ecological trait. Reported are the estimated correlation parameter ($\lambda$, $\alpha$, or $a$), log-likelihood (logLik), number of parameters ($k$), standard error (se), expected information criterion (EIC), ΔEIC, and model weight. For ecological predictors, permutation MANOVA results are provided using Pillai's trace with 999 permutations. Models with ΔEIC $< 2$ or weight $> 0.1$ are highlighted in bold as best supported.

| Predictor | Model | $\lambda/\alpha/a$ | logLik | k | se | EIC | ΔEIC | $EIC_w$ | Pillai | $p$-value |
|---|---|---|---|---|---|---|---|---|---|---|
| **Global** | | | | | | | | | | |
| ~1 | BM | — | -26472.552 | 16 | 14.891 | 53731.302 | 5029.777 | 0.000 | — | — |
| ~1 | Pagel's $\lambda$ | 0.500 | -24174.770 | 16 | 3.137 | 48718.253 | 16.728 | 0.000 | — | — |
| ~1 | OU | 0.715 | -24628.180 | 16 | 3.244 | 49618.060 | 916.535 | 0.000 | — | — |
| ~1 | EB | 0.000 | -26472.535 | 16 | 16.837 | 53769.479 | 5067.954 | 0.000 | — | — |
| ~Diet | BM | — | -26379.608 | 16 | 13.436 | 53510.300 | 4808.775 | 0.000 | 0.411 | 0.001 |
| **~Diet** | **Pagel's $\lambda$** | **0.500** | **-24131.917** | **16** | **3.213** | **48701.525** | **0.000** | **0.860** | **0.225** | 0.001 |
| ~Diet | OU | 0.750 | -24521.072 | 16 | 3.602 | 49475.418 | 773.893 | 0.000 | 0.481 | 0.001 |
| ~Diet | EB | 0.000 | -26383.862 | 16 | 17.399 | 53672.915 | 4971.390 | 0.000 | 0.410 | 0.001 |
| ~Habitat | BM | — | -26347.680 | 16 | 13.857 | 53491.733 | 4790.208 | 0.000 | 0.573 | 0.006 |
| ~Habitat | Pagel's $\lambda$ | 0.500 | -24124.024 | 16 | 4.013 | 48745.559 | 44.034 | 0.000 | 0.284 | 0.002 |
| ~Habitat | OU | 0.974 | -24443.482 | 16 | 4.082 | 49394.321 | 692.796 | 0.000 | 0.804 | 0.001 |
| ~Habitat | EB | 0.000 | -26350.980 | 16 | 17.299 | 53659.618 | 4958.093 | 0.000 | 0.572 | 0.004 |
| ~Symbiosis | BM | — | -26400.128 | 16 | 14.277 | 53606.339 | 4904.814 | 0.000 | 0.359 | 0.009 |
| **~Symbiosis** | **Pagel's $\lambda$** | **0.500** | **-24136.555** | **16** | **3.077** | **48705.153** | **3.628** | **0.140** | **0.208** | 0.002 |
| ~Symbiosis | OU | 0.920 | -24460.985 | 16 | 3.518 | 49357.552 | 656.028 | 0.000 | 0.704 | 0.001 |
| ~Symbiosis | EB | 0.000 | -26403.103 | 16 | 17.360 | 53685.585 | 4984.061 | 0.000 | 0.359 | 0.015 |
| **Regime-dependent** | | | | | | | | | | |
| ~1 | BMM\| Diet | — | -26293.531 | 16 | 11.062 | 53317.405 | 4615.880 | 0.000 | — | — |
| ~1 | BMM\| Habitat | — | -26224.460 | 16 | 11.972 | 53160.780 | 4459.256 | 0.000 | — | — |
| ~1 | BMM\| Symbiosis | — | -27448.374 | 16 | 18.966 | 55674.761 | 6973.236 | 0.000 | — | — |
| ~1 | OUM\| Diet | 0.665 | -24520.956 | 16 | 3.939 | 49497.730 | 796.206 | 0.000 | — | — |
| ~1 | OUM\| Habitat | 0.721 | -24434.451 | 16 | 3.936 | 49390.721 | 689.197 | 0.000 | — | — |
| ~1 | OUM\| Symbiosis | 0.820 | -24459.123 | 16 | 3.257 | 49359.199 | 657.674 | 0.000 | — | — |

**Table 10**: Model comparison of multivariate phylogenetic GLS (mvGLS) analyses testing ecological predictors of morphology principal components (first 8 PCs). Global models (BM, Pagel's $\lambda$, OU, EB) were fitted on the phylogeny, while regime-dependent models (BMM: multiple-rate BM; OUM: multiple-optima OU) were fitted on a consensus stochastic character map (SCM) for each ecological trait. Reported are the estimated correlation parameter ($\lambda$, $\alpha$, or $a$), log-likelihood (logLik), number of parameters ($k$), standard error (se), expected information criterion (EIC), ΔEIC, and model weight. For ecological predictors, permutation MANOVA results are provided using Pillai's trace with 999 permutations. Models with ΔEIC $< 2$ or weight $> 0.1$ are highlighted in bold as best supported.

| Predictor | Model | $\lambda/\alpha/a$ | logLik | k | se | EIC | ΔEIC | $EIC_w$ | Pillai | $p$-value |
|---|---|---|---|---|---|---|---|---|---|---|
| **Global** | | | | | | | | | | |
| ~1 | BM | — | -12635.456 | 11 | 8.571 | 25686.428 | 3527.141 | 0.000 | — | — |
| ~1 | Pagel's $\lambda$ | 0.603 | -10987.340 | 11 | 2.453 | 22186.205 | 26.918 | 0.000 | — | — |
| ~1 | OU | 0.913 | -11426.927 | 11 | 2.505 | 23045.228 | 885.941 | 0.000 | — | — |
| ~1 | EB | 0.000 | -12635.457 | 11 | 12.920 | 25785.579 | 3626.292 | 0.000 | — | — |
| ~Diet | BM | — | -12598.845 | 11 | 8.321 | 25651.294 | 3492.007 | 0.000 | 0.206 | 0.011* |
| **~Diet** | **Pagel's $\lambda$** | **0.579** | **-10954.419** | **11** | **2.637** | **22159.287** | **0.000** | **1.000** | **0.177** | **0.001*** |
| ~Diet | OU | 1.053 | -11321.140 | 11 | 2.058 | 22878.782 | 719.495 | 0.000 | 0.466 | 0.001* |
| ~Diet | EB | 0.000 | -12598.851 | 11 | 15.821 | 25740.259 | 3580.972 | 0.000 | 0.206 | 0.010* |
| ~Habitat | BM | — | -12570.147 | 11 | 8.301 | 25605.532 | 3446.246 | 0.000 | 0.338 | 0.019* |
| ~Habitat | Pagel's $\lambda$ | 0.599 | -10960.030 | 11 | 2.527 | 22209.297 | 50.010 | 0.000 | 0.157 | 0.145 |
| ~Habitat | OU | 1.412 | -11306.720 | 11 | 2.320 | 22887.691 | 728.404 | 0.000 | 0.578 | 0.001* |
| ~Habitat | EB | 0.000 | -12570.150 | 11 | 13.183 | 25737.512 | 3578.225 | 0.000 | 0.338 | 0.010* |
| ~Symbiosis | BM | — | -12616.785 | 11 | 9.094 | 25682.916 | 3523.629 | 0.000 | 0.114 | 0.099 |
| ~Symbiosis | Pagel's $\lambda$ | 0.600 | -10962.622 | 11 | 2.461 | 22180.347 | 21.060 | 0.000 | 0.136 | 0.006* |
| ~Symbiosis | OU | 1.303 | -11310.690 | 11 | 2.476 | 22859.828 | 700.541 | 0.000 | 0.515 | 0.001* |
| ~Symbiosis | EB | 0.000 | -12616.788 | 11 | 15.266 | 25805.207 | 3645.920 | 0.000 | 0.114 | 0.105 |
| **Regime-dependent** | | | | | | | | | | |
| ~1 | BMM\| Diet | — | -12583.386 | 11 | 11.630 | 25666.826 | 3507.539 | 0.000 | — | — |
| ~1 | BMM\| Habitat | — | -12440.250 | 11 | 7.068 | 25329.162 | 3169.875 | 0.000 | — | — |
| ~1 | BMM\| Symbiosis | — | -12478.051 | 11 | 6.801 | 25362.830 | 3203.543 | 0.000 | — | — |
| ~1 | OUM\| Diet | 0.791 | -11317.433 | 11 | 2.692 | 22883.360 | 724.073 | 0.000 | — | — |
| ~1 | OUM\| Habitat | 0.808 | -11305.635 | 11 | 2.490 | 22919.173 | 759.886 | 0.000 | — | — |
| ~1 | OUM\| Symbiosis | 1.025 | -11310.328 | 11 | 2.228 | 22849.905 | 690.618 | 0.000 | — | — |

**Table 11**: Model comparison of multivariate phylogenetic GLS (mvGLS) analyses testing ecological predictors of colour principal components (first 8 PCs). Global models (BM, Pagel's $\lambda$, OU, EB) were fitted on the phylogeny, while regime-dependent models (BMM: multiple-rate BM; OUM: multiple-optima OU) were fitted on a consensus stochastic character map (SCM) for each ecological trait. Reported are the estimated correlation parameter ($\lambda$, $\alpha$, or $a$), log-likelihood (logLik), number of parameters ($k$), standard error (se), expected information criterion (EIC), ΔEIC, and model weight. For ecological predictors, permutation MANOVA results are provided using Pillai's trace with 999 permutations. Models with ΔEIC $< 2$ or weight $> 0.1$ are highlighted in bold as best supported.

| Predictor | Model | $\lambda/\alpha/a$ | logLik | k | se | EIC | ΔEIC | $EIC_w$ | Pillai | $p$-value |
|---|---|---|---|---|---|---|---|---|---|---|
| **Global** | | | | | | | | | | |
| ~1 | BM | — | -16681.545 | 8 | 4.334 | 33549.301 | 2505.581 | 0.000 | — | — |
| ~1 | Pagel's $\lambda$ | 0.514 | -15474.040 | 8 | 1.694 | 31063.457 | 19.737 | 0.000 | — | — |
| ~1 | OU | 0.576 | -15608.744 | 8 | 2.113 | 31334.682 | 290.962 | 0.000 | — | — |
| ~1 | EB | 0.000 | -16681.545 | 8 | 7.446 | 33620.466 | 2576.746 | 0.000 | — | — |
| ~Diet | BM | — | -16640.782 | 8 | 4.458 | 33498.114 | 2454.394 | 0.000 | 0.203 | 0.002* |
| ~Diet | Pagel's $\lambda$ | 0.508 | -15461.268 | 8 | 1.795 | 31065.210 | 21.490 | 0.000 | 0.072 | 0.050 |
| ~Diet | OU | 0.585 | -15578.011 | 8 | 1.630 | 31296.376 | 252.656 | 0.000 | 0.167 | 0.001* |
| ~Diet | EB | 0.000 | -16640.782 | 8 | 8.283 | 33566.835 | 2523.116 | 0.000 | 0.203 | 0.003* |
| ~Habitat | BM | — | -16604.242 | 8 | 4.337 | 33457.792 | 2414.072 | 0.000 | 0.368 | 0.005* |
| ~Habitat | Pagel's $\lambda$ | 0.478 | -15445.195 | 8 | 1.916 | 31067.436 | 23.717 | 0.000 | 0.160 | 0.012* |
| ~Habitat | OU | 0.801 | -15511.066 | 8 | 2.085 | 31195.654 | 151.935 | 0.000 | 0.463 | 0.001* |
| ~Habitat | EB | 0.000 | -16604.242 | 8 | 8.061 | 33534.436 | 2490.716 | 0.000 | 0.368 | 0.004* |
| ~Symbiosis | BM | — | -16631.451 | 8 | 4.750 | 33487.916 | 2444.196 | 0.000 | 0.251 | 0.005* |
| **~Symbiosis** | **Pagel's $\lambda$** | **0.483** | **-15453.571** | **8** | **1.772** | **31043.720** | **0.000** | **1.000** | **0.112** | **0.005*** |
| ~Symbiosis | OU | 0.814 | -15504.776 | 8 | 1.918 | 31147.578 | 103.858 | 0.000 | 0.487 | 0.001* |
| ~Symbiosis | EB | 0.000 | -16631.451 | 8 | 7.529 | 33553.186 | 2509.466 | 0.000 | 0.251 | 0.006* |
| **Regime-dependent** | | | | | | | | | | |
| ~1 | BMM\| Diet | — | -16530.313 | 8 | 5.270 | 33284.409 | 2240.689 | 0.000 | — | — |
| ~1 | BMM\| Habitat | — | -16457.505 | 8 | 4.514 | 33136.452 | 2092.733 | 0.000 | — | — |
| ~1 | BMM\| Symbiosis | — | -16467.154 | 8 | 3.857 | 33149.102 | 2105.382 | 0.000 | — | — |
| ~1 | OUM\| Diet | 0.547 | -15578.828 | 8 | 1.878 | 31305.289 | 261.569 | 0.000 | — | — |
| ~1 | OUM\| Habitat | 0.692 | -15508.576 | 8 | 2.272 | 31211.019 | 167.299 | 0.000 | — | — |
| ~1 | OUM\| Symbiosis | 0.750 | -15503.266 | 8 | 1.969 | 31156.609 | 112.889 | 0.000 | — | — |

**Table 12**: Phylogenetic generalized least squares (PGLS) associations between colour and morphological principal components (PCs). Shown are the PC pairing, slope, raw *p*-value, coefficient of determination ($R^2$), best-fitting correlation structure (Pagel or OU), and FDR-adjusted *p*-value.

| Colour | Morphology | Best model | $R^2$ | Slope | p | p(FDR) |
|---|---|---|---|---|---|---|
| PC1 | PC1 | Pagel | 0.001 | -0.073 | 0.2440 | 0.3828 |
| PC1 | PC2 | Pagel | 0.006 | -0.061 | 0.2804 | 0.3828 |
| PC1 | PC3 | Pagel | 0.008 | 0.026 | 0.6653 | 0.7604 |
| PC1 | PC4 | Pagel | 0.022 | 0.063 | 0.2871 | 0.3828 |
| PC1 | PC5 | Pagel | 0.021 | -0.135 | 0.0156 | 0.1249 |
| PC1 | PC6 | Pagel | 0.021 | -0.126 | 0.0981 | 0.3828 |
| PC1 | PC7 | Pagel | 0.001 | -0.085 | 0.1646 | 0.3828 |
| PC1 | PC8 | Pagel | 0.000 | 0.014 | 0.7916 | 0.7916 |
| PC2 | PC1 | Pagel | 0.008 | -0.104 | 0.0603 | 0.2926 |
| PC2 | PC2 | Pagel | 0.003 | -0.047 | 0.3393 | 0.4263 |
| PC2 | PC3 | Pagel | 0.005 | 0.047 | 0.3730 | 0.4263 |
| PC2 | PC4 | Pagel | 0.010 | 0.066 | 0.2025 | 0.3404 |
| PC2 | PC5 | Pagel | 0.021 | 0.008 | 0.8759 | 0.8759 |
| PC2 | PC6 | Pagel | 0.000 | 0.108 | 0.1097 | 0.2926 |
| PC2 | PC7 | Pagel | 0.002 | -0.096 | 0.0733 | 0.2926 |
| PC2 | PC8 | Pagel | 0.023 | 0.058 | 0.2128 | 0.3404 |
| PC3 | PC1 | Pagel | 0.022 | 0.097 | 0.0993 | 0.1987 |
| **PC3** | **PC2** | **Pagel** | **0.020** | **0.158** | **0.0051** | **0.0409*** |
| PC3 | PC3 | Pagel | 0.009 | -0.112 | 0.0656 | 0.1748 |
| PC3 | PC4 | Pagel | 0.007 | 0.040 | 0.4975 | 0.5772 |
| PC3 | PC5 | Pagel | 0.005 | 0.066 | 0.2396 | 0.3834 |
| PC3 | PC6 | Pagel | 0.016 | -0.048 | 0.5051 | 0.5772 |
| PC3 | PC7 | Pagel | 0.000 | 0.028 | 0.6481 | 0.6481 |
| PC3 | PC8 | Pagel | 0.020 | 0.131 | 0.0171 | 0.0683 |
| PC4 | PC1 | Pagel | 0.064 | 0.142 | 0.0114 | 0.0914 |
| PC4 | PC2 | Pagel | 0.007 | 0.080 | 0.1092 | 0.1747 |
| PC4 | PC3 | Pagel | 0.000 | -0.016 | 0.7678 | 0.7678 |
| PC4 | PC4 | Pagel | 0.009 | 0.030 | 0.5624 | 0.6427 |
| PC4 | PC5 | Pagel | 0.000 | -0.050 | 0.3123 | 0.4164 |
| PC4 | PC6 | Pagel | 0.022 | 0.117 | 0.0890 | 0.1747 |
| PC4 | PC7 | Pagel | 0.001 | -0.100 | 0.0681 | 0.1747 |
| PC4 | PC8 | Pagel | 0.006 | 0.078 | 0.0988 | 0.1747 |
| **PC5** | **PC1** | **Pagel** | **0.027** | **0.167** | **0.0052** | **0.0352*** |
| PC5 | PC2 | Pagel | 0.003 | -0.005 | 0.9235 | 0.9941 |
| PC5 | PC3 | Pagel | 0.004 | -0.000 | 0.9941 | 0.9941 |
| PC5 | PC4 | Pagel | 0.002 | 0.021 | 0.7217 | 0.9941 |
| PC5 | PC5 | Pagel | 0.002 | -0.006 | 0.9165 | 0.9941 |
| **PC5** | **PC6** | **OU** | **0.020** | **-0.143** | **0.0088** | **0.0352*** |
| PC5 | PC7 | Pagel | 0.006 | -0.070 | 0.2460 | 0.4920 |
| PC5 | PC8 | Pagel | 0.012 | 0.095 | 0.0828 | 0.2209 |
| PC6 | PC1 | Pagel | 0.000 | 0.050 | 0.3951 | 0.4516 |
| PC6 | PC2 | Pagel | 0.003 | -0.049 | 0.3440 | 0.4516 |
| PC6 | PC3 | Pagel | 0.001 | 0.058 | 0.3067 | 0.4516 |
| **PC6** | **PC4** | **Pagel** | **0.045** | **-0.161** | **0.0034** | **0.0271*** |
| PC6 | PC5 | Pagel | 0.003 | -0.104 | 0.0439 | 0.1756 |
| PC6 | PC6 | Pagel | 0.070 | 0.095 | 0.1864 | 0.3728 |
| PC6 | PC7 | Pagel | 0.005 | 0.079 | 0.1664 | 0.3728 |
| PC6 | PC8 | Pagel | 0.009 | 0.033 | 0.5014 | 0.5014 |
| PC7 | PC1 | Pagel | 0.000 | -0.112 | 0.0633 | 0.1267 |
| PC7 | PC2 | Pagel | 0.007 | -0.102 | 0.0500 | 0.1267 |
| **PC7** | **PC3** | **Pagel** | **0.045** | **0.177** | **0.0016** | **0.0132*** |
| PC7 | PC4 | Pagel | 0.024 | 0.090 | 0.1005 | 0.1608 |
| PC7 | PC5 | Pagel | 0.000 | -0.006 | 0.9005 | 0.9005 |
| **PC7** | **PC6** | **Pagel** | **0.004** | **-0.185** | **0.0116** | **0.0464*** |
| PC7 | PC7 | Pagel | 0.003 | -0.023 | 0.6968 | 0.9005 |
| PC7 | PC8 | Pagel | 0.003 | -0.007 | 0.8864 | 0.9005 |
| PC8 | PC1 | OU | 0.012 | -0.117 | 0.0330 | 0.1977 |
| PC8 | PC2 | OU | 0.001 | -0.034 | 0.5318 | 0.8319 |
| PC8 | PC3 | OU | 0.000 | 0.014 | 0.7974 | 0.8319 |
| PC8 | PC4 | OU | 0.000 | -0.015 | 0.7851 | 0.8319 |
| PC8 | PC5 | OU | 0.000 | 0.011 | 0.8319 | 0.8319 |
| PC8 | PC6 | OU | 0.013 | -0.107 | 0.0494 | 0.1977 |
| PC8 | PC7 | OU | 0.010 | -0.091 | 0.0956 | 0.2550 |
| PC8 | PC8 | OU | 0.001 | 0.013 | 0.8094 | 0.8319 |

**Table 13**: Multivariate PGLS models testing the effect of morphology on colour patterns across Pomacentridae. Models compared using the Expected Information Criterion (EIC). Parameter estimates ($\lambda$, $\alpha$, $a$) are reported when applicable. Pillai's trace statistic and permutation *p*-value are shown for the best supported trait model.

| model | $\lambda/\alpha/a$ | predictor | logLik | k | se | EIC | ΔEIC | weight | Pillai | *p*-value |
|---|---|---|---|---|---|---|---|---|---|---|
| BM | — | ~1 | -16681.55 | 8 | 3.777 | 33547.30 | 2495.831 | 0.000 | — | — |
| BM | — | ~morphology | -16479.01 | 8 | 2.738 | 33221.59 | 2170.123 | 0.000 | — | — |
| Pagel's $\lambda$ | 0.513 | ~1 | -15474.04 | 8 | 1.553 | 31062.24 | 10.769 | 0.456 | — | — |
| **Pagel's $\lambda$** | **0.497** | **~morphology** | **-15408.34** | **8** | **2.134** | **31051.47** | **0.00** | **0.995** | **0.356** | **0.001*** |
| OU | 0.575 | ~1 | -15608.74 | 8 | 1.769 | 31330.99 | 279.518 | 0.000 | — | — |
| OU | 0.707 | ~morphology | -15499.62 | 8 | 2.163 | 31234.41 | 182.940 | 0.000 | — | — |
| EB | 0.000 | ~1 | -16681.54 | 8 | 7.483 | 33619.75 | 2568.284 | 0.000 | — | — |
| EB | 0.000 | ~morphology | -16479.01 | 8 | 5.073 | 33293.28 | 2241.812 | 0.000 | — | — |
